# Loss of STAT5 in adipocytes increases subcutaneous fat mass via sex-dependent and depot-specific pathways

**DOI:** 10.1101/2021.03.24.436884

**Authors:** Allison J. Richard, Hardy Hang, Timothy D. Allerton, Peng Zhao, Sujoy Ghosh, Carrie M. Elks, Jacqueline M. Stephens

## Abstract

The STAT (Signal Transducers and Activators of Transcription) family of transcription factors contributes to adipocyte development and function. STAT5A and STAT5B are induced during adipocyte differentiation and are primarily activated by growth hormone (GH). Studies in mice lacking adipocyte GH receptor or STAT5 support their roles in lipolysis-mediated reduction of adipose tissue mass. We have generated a mouse model lacking both STAT5 genes specifically in adipocytes (STAT5^AKO^). Notably, both sexes of STAT5^AKO^ mice have increased inguinal adipose tissue without any changes in gonadal fat mass. However, both depots exhibit substantial differences in fat cell size. Study of STAT5^AKO^ mice also have revealed that GH’s ability to induce insulin resistance is dependent upon STAT5 in adipocytes, but its ability to reduce adipose tissue mass is STAT5 independent. Additional observations, which were not predicted, indicate that the causes and regulation of increased fat mass in STAT5^AKO^ mice are sex- and depot-dependent.

## INTRODUCTION

Adipocytes have long been recognized as storage sites for excess energy. It is now well recognized that fat cells are insulin sensitive, have endocrine functions, and serve as important regulators of whole-body energy homeostasis (Sethi and Vidal-Puig, 2007). Obesity is the primary disease of adipocytes and is a prominent risk factor for the development of type 2 diabetes mellitus (T2DM), cardiovascular disease, and certain cancers. However, some people with obesity are metabolically healthy (Stefan et al., 2018) and it is well-established that subcutaneous fat can have protective roles against metabolic dysfunction (Booth et al., 2018). Mouse models and humans with reduced growth hormone (GH) signaling have increased subcutaneous adiposity but improved metabolic health (Berryman et al., 2004; Coschigano et al., 2003; Dominici et al., 2002; Guevara-Aguirre et al., 2020a, 2020b).

The Janus kinase/signal transducer and activator of transcription (JAK/STAT) pathway is used to transmit cellular signaling of many hormones and cytokines (Pellegrini and Dusanter-Fourt, 1997). JAKs utilize plasma membrane receptors to activate signal transduction cascades involving STAT proteins (Morris et al., 2018). Seven proteins comprise the STAT family of mammalian transcription factors (STATs 1, 2, 3, 4, 5A, 5B, and 6), which, in response to receptor stimulation, are phosphorylated on tyrosine residues causing their nuclear translocation. Each STAT has a distinct pattern of activation. Upon nuclear translocation, STATs bind distinct DNA sequences to regulate transcription (Darnell, 1997; Pellegrini and Dusanter-Fourt, 1997). The specificity of STAT activation and function is still not completely understood, but is determined, at least in part, by the receptor and the specific STAT protein. Two STAT family members, STAT5A and STAT5B, are highly related, but are encoded for by separate genes (Liu et al., 1995). Murine STAT5A and 5B have 96% sequence similarity and both STAT5 isoforms can have essential and nonessential, or redundant, roles in hormone responses (Kornfeld et al., 2008; Teglund et al., 1998). STAT5 proteins have been reported to be activated by a number of different cytokines/hormones, but the phenotype of STAT5-null mice is consistent with STAT5 proteins being primarily activated by growth hormone and prolactin (Teglund et al., 1998). Polymorphisms in STAT5 genes are primarily associated with different types of cancers or changes in milk production in a variety of species. However, STAT5 polymorphisms are also associated with altered cholesterol metabolism (Makimura et al., 2011), body weight (Zhao et al., 2012), and lipid metabolism (Meirhaeghe et al., 2003). The JAK2-STAT5 signaling pathway contributes to obesity pathogenesis and T2DM by impacting a variety of different tissues (Gurzov et al., 2016). Moreover, there is compelling data to show disruption of the JAK2-STAT5 pathway in adipocytes (Kaltenecker et al., 2017) can improve both systemic metabolic functions and liver function (Corbit et al., 2017a, 2018, 2019).

There are many studies that show that STAT5 proteins regulate adipocyte development (Floyd and Stephens, 2003; Jung et al., 2012; Kawai et al., 2007; Liu et al., 2020; Miyaoka et al., 2006; Nanbu-Wakao et al., 2002; Richter et al., 2003; Shang and Waters, 2003; Stewart et al., 2004, 2011; Wakao et al., 2011; Wang et al., 2018; Yao et al., 2019; Yarwood et al., 1999), fat mass (Teglund et al., 1998) and lipid metabolism (Meirhaeghe et al., 2003). Yet, the specific mechanisms involved remain largely unknown. To identify the functions of adipocyte STAT5 proteins, we generated mice lacking both STAT5 genes in adipocytes (STAT5^AKO^). STAT5^AKO^ mice have prominent increased subcutaneous adiposity on chow diet and have indicators of improved insulin sensitivity. Notably, other features of these mice were not predicted and have provided observations related to lipid metabolism and GH actions on adipocytes that challenge the current dogma. These observations suggest there are critical knowledge gaps regarding adipocyte STAT5 and its ability to contribute to systemic metabolic regulation. It is largely accepted that GH signaling reduces fat mass by promoting lipolysis and STAT5 is the primary mediator of GH action. However, characterization of STAT5^AKO^ mice reveals that the GH-induced loss of fat mass occurs in a STAT5-independent manner. We also observe that the increased adiposity of STAT5^AKO^ mice is *not* associated with changes in adipose tissue lipolysis. Our data, and studies from other groups (Corbit et al., 2017a, 2018, 2019), strongly indicate that the mechanism(s) by which the adipocyte JAK2-STAT5 pathway contributes to systemic metabolism are more complex than originally thought. Overall, our studies show that congenital loss of STAT5 in mature adipocytes results in a depot-specific increase in fat mass as well as sex-dependent differences in adipocyte gene expression and whole-body energy expenditure.

## RESULTS

### Generation of adipocyte-specific STAT5A and STAT5B knockout (STAT5^AKO^) mice

Mice with both the *Stat5a* and *Stat5b* genes flanked by a set of loxP sites (floxed) were bred to adiponectin-Cre (AdipoQ-Cre) mice to produce mice with a tissue-specific double knockout of both STATs 5a & 5b (Cui et al., 2004). Both mouse strains have been maintained on a C57BL/6J background. STAT5^fl/+:AdipoQ-Cre/+^ mice were crossed with STAT5^fl/fl:+/+^ mice to create STAT5^AKO^ mice (STAT5^fl/fl:AdipoQ-Cre/+^) and control littermates (STAT5^fl/fl:+/+^). Genotype data of female mice is shown in Figure 1. Both STAT 5A and B proteins (Figures 1A and 1B) and mRNA (Figures 1C and 1D) were reduced, but not absent, from fat tissue homogenates prepared from STAT5^AKO^ mice. This was expected since STAT5 is also expressed in non-adipocyte cells. We also examined the expression of STAT5 mRNA in several fat pads as well as liver and skeletal muscle (Figures 1C and 1D). *Stat5a* mRNA was reduced about 50% in iWAT and gonadal fat (gWAT) and there was greater than a 60% reduction of *Stats 5a* and *5b* mRNA in brown adipose tissue (BAT). There were no changes in *Stat5* gene expression in liver or skeletal muscle. Growth hormone (GH) rapidly induces *Cish* gene expression in a STAT5-dependent manner (Clasen et al., 2013). Whereas acute GH injection increased *Cish* mRNA expression in the liver of both floxed and STAT5^AKO^ mice, in the iWAT it only induced *Cish* mRNA in floxed, but not STAT5^AKO^ mice (Figure 1E). Since STAT5 proteins are also expressed in cells present in the stromal vascular fraction (SVF) of adipose tissue (AT), we fractionated fat to assess knockdown efficiency. As expected, we observed a decrease in STAT5 proteins in AKO heterozygotes (het) and a substantial (>85%) reduction in STAT5A and STAT5B in the adipocyte fraction from STAT5^AKO^ mice (Figure 1F). We also observe a consistent increase of STAT5A protein in the SVF cell population of STAT5^AKO^ mice. There were no changes in STAT5 protein levels in liver or skeletal muscle in STAT5^AKO^ female mice (data not shown). We observed similar observations in the genotypic analysis of male mice (Figure S1). Collectively, our results demonstrate Cre-mediated excision of *STAT5a/b* only in mature adipocytes of male and female mice.

**Figure 1.**
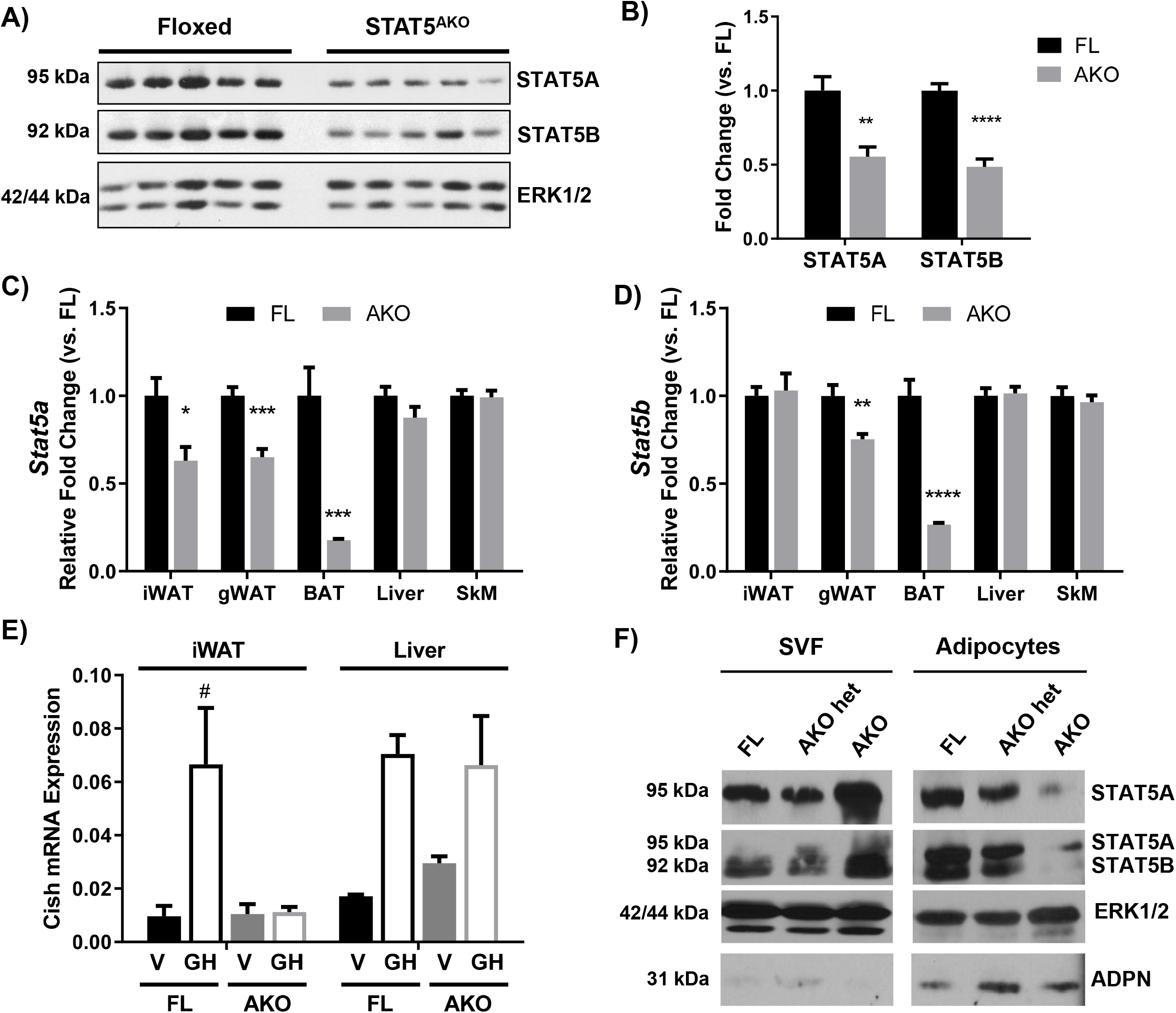
Expression of STAT5A and 5B is knocked down in adipose tissue and adipocytes of STAT5^AKO^ female mice. Female STAT5^AKO^ (AKO) mice and their floxed (FL) littermate controls were euthanized at 3 (A-D), 11 (E), or 4 (F) months of age and tissues were immediately collected for protein or gene expression analyses (A-E) or cellular fractionation (F). A) Immunoblot of proteins resolved from iWAT samples for 5 individual mice of each genotype. B) Quantification of band intensities from A (n = 5 per group). ERK1/2 was used as a loading control, and band intensities were normalized to ERK1/2 and then represented as fold change relative to FL mice. C and D) *Stat5a* and *Stat5b* gene expression measured by RT-qPCR for the indicated tissues (n = 5 – 8 mice per group). E) Eleven-month-old female mice were injected with 1.5mg/kg mGH or vehicle (V; 0.1% BSA/PBS) for 30 minutes prior to euthanasia and tissue collection. Gene expression measured by RT-qPCR is shown (n = 2 – 3 mice per group). F) Gonadal white adipose tissue was removed from female STAT5^AKO^, heterozygous STAT5^AKO^ (het AKO), and homozygous floxed control (FL) mice, fractionated into adipocytes and stromal vascular fraction (SVF), and 100 μg (SVF) or 200 μg (adipocyte) protein subjected to western blotting. Also shown are ERK1/2 as a loading control and adiponectin (ADPN) as an adipocyte marker. For each protein, the SVF and adipocyte samples were run on the same gel and the images were from the same exposure of the same blot. Significance was determined by *t*-test for FL versus AKO comparisons in A-D) and is denoted as * p < 0.05, ** p < 0.01, *** p < 0.001, **** p < 0.0001. For E, a 2-way ANOVA was used to assess treatment/genotype and tissue variables with Tukey’s post-hoc multiple comparisons test to compare all treatment/genotype groups for each tissue; # denotes p < 0.05 for GH versus V comparisons. See also Figure S1.

### Depot-specific differences in adipose tissue mass and adipocyte size in STAT5^AKO^ mice

As shown in Figures 2A and 2B, female STAT5^AKO^ mice have a significant increase in inguinal WAT weight, without having discernable changes in gonadal WAT or other fat depots with exception of statistically significant increase in retroperitoneal and brown fat. Notably, only the subcutaneous iWAT depot of STAT5^AKO^ mice was consistently increased when several cohorts of mice of different ages were compared (data not shown). STAT5^AKO^ female mice do not have altered serum IGF-1 levels (Figure 2D), but there was a trend towards decreased GH levels in STAT5^AKO^ females (Figure 2C). Notably, these mice have a significant decrease in fasting insulin levels, but not glucose levels (Figures 2E and 2F). These results reveal that STAT5^AKO^ females have increased adiposity and may have improved insulin sensitivity on chow or low-fat diets as indicated by HOMA-IR (Figure 2G).

**Figure 2.**
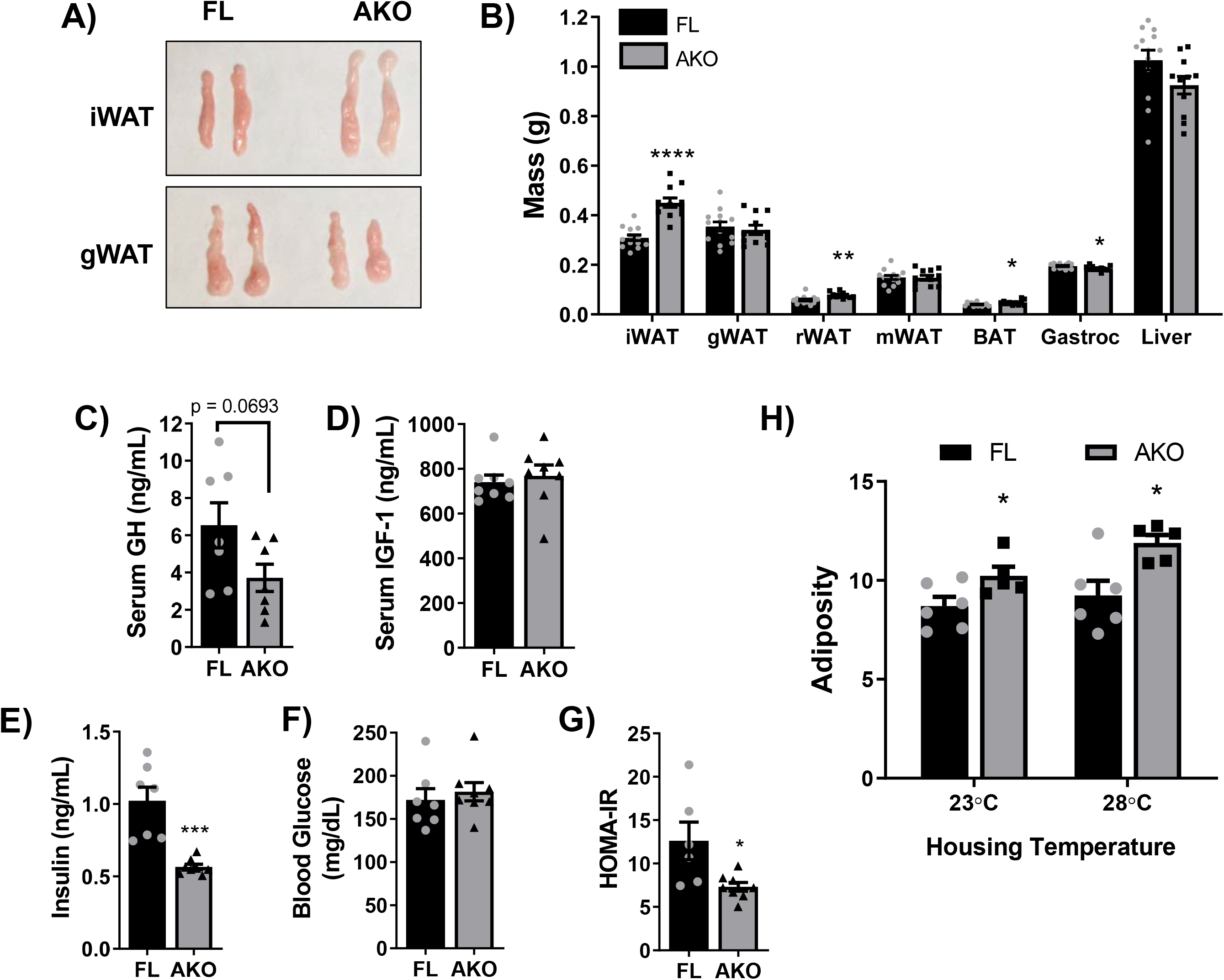
Female STAT5^AKO^ mice have increased adiposity and are metabolically healthier than floxed controls when fed chow or low-fat diet. Female STAT5^AKO^ (AKO) and floxed (FL) littermate control mice were weaned onto regular chow diet (13% kcal from fat) and maintained on that diet (A, B, and H) or switched to a defined-composition low fat diet (LFD - 10% kcal from fat; C - G) at 6 weeks of age. A) Representative images of inguinal and gonadal white adipose tissue depots (iWAT and gWAT) collected from five-month-old mice. B) Tissue weights of white adipose tissue depots (iWAT, gWAT, retroperitoneal - rWAT, mesenteric - mWAT), brown adipose tissue (BAT), gastrocnemius skeletal muscle (Gastroc), and liver collected from 10-week-old mice (n = 11-13). C – F) Serum growth hormone (GH), insulin growth factor 1 (IGF-1), insulin, and blood glucose levels collected from 3-month-old mice on LFD for 1 month (n = 7 – 8). G) HOMA-IR was calculated from insulin and glucose levels in E and F. H) Mice (n = 5-6 per group) were housed at different temperatures beginning at weaning (3 weeks of age). They were fed chow diet (13% kcal from fat) for 6 weeks and then body weight and body composition (fat mass and lean mass) were measured at 9 weeks of age. Adiposity was calculated as fat mass divided by total body weight for each animal. Significance was determined by *t*-test and is denoted as * p < 0.05, ** p < 0.01, *** p < 0.001, or **** p < 0.0001 for AKO versus FL comparisons. See also Figure S2.

We observed similar, but not identical, outcomes in male STAT5^AKO^ mice (Figure S2). Like females, STAT5^AKO^ male mice had increased subcutaneous fat mass and improved HOMA-IR, but distinct from the females, the decreased HOMA-IR in STAT5^AKO^ male mice was driven by lower glucose values instead of reduced insulin levels. The range of circulating GH and IGF-1 levels was larger, but these values did not significantly differ between genotypes (Figure S2). In many mouse models, alterations in fat mass are negated at thermoneutrality. To assess the robustness of the adiposity phenotype in STAT5^AKO^ mice, we also examined body composition at thermoneutrality. As shown in Figures 2H and S2H, both male and female mice maintained increased adiposity following six weeks of thermoneutral housing conditions.

Increased adipose tissue mass is typically associated with hypertrophy and/or hyperplasia. Histological analysis reveals hypertrophic adipocytes in both iWAT and gWAT of female (Figure 3) and male STAT5^AKO^ mice (Figure S3) that is accompanied by a decrease in the number of smaller adipocytes.

**Figure 3.**
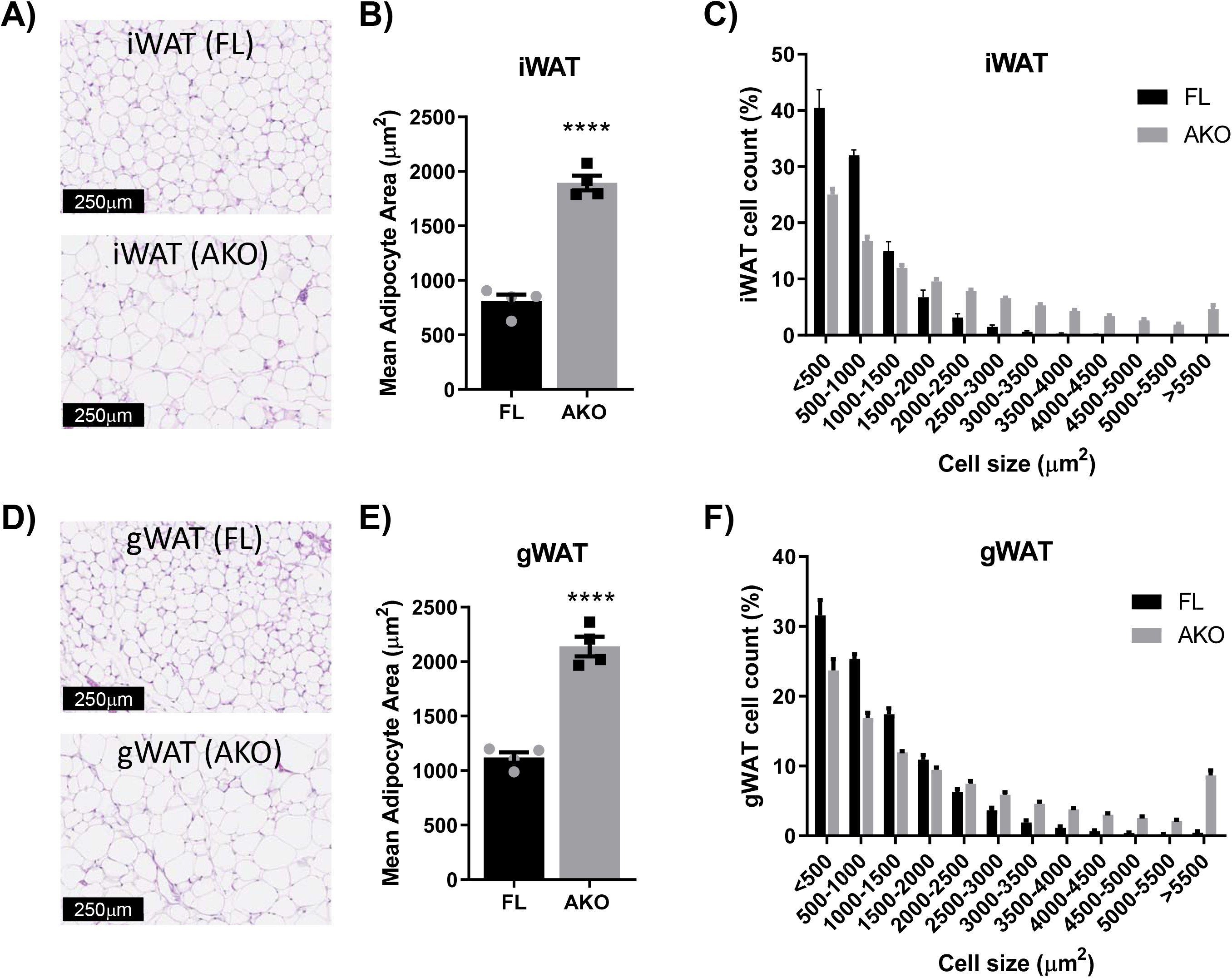
Female STAT5^AKO^ mice have larger fat cells than floxed control mice in both subcutaneous and visceral adipose tissue depots when fed a chow diet. Female STAT5^AKO^ (AKO) and floxed (FL) littermate control mice were weaned onto regular chow diet (13% kcal from fat). A and D) Representative images of H&E stained inguinal and gonadal white adipose tissue depots (iWAT and gWAT) collected from five-month-old mice. Quantification of total mean adipocyte area (B and E) and fat cell size distribution (C and F) from H&E-stained images as shown in A and D (n = 4 mice per genotype). Significance in B and E was determined by *t*-test and is denoted as **** p < 0.0001 for AKO versus FL comparisons. See also Figure S3.

### Lipid Metabolism and lipolytic responses in STAT5^AKO^ mice

It is largely accepted that GH promotes fat loss via lipolysis and animal models with diminished adipose tissue GH signaling have altered lipolytic responses (Corbit et al., 2017b, 2018, 2019; Kaltenecker et al., 2017; List et al., 2013; Nordstrom et al., 2013). We examined gene and protein expression of several known lipolytic mediators including adipose triglyceride lipase (ATGL), CGI-58 (comparative gene identification-58), and β3 adrenergic receptor (ADRB3). As shown in Figure S4, male STAT5^AKO^ mice have decreased gene expression of *Atgl/Pnpla2, Cgi-58/Abhd5* and *Adrb3*, but reduced gene expression only translates to diminished protein expression levels for CGI-58. Circulating levels of glycerol, triglycerides (TG), and free fatty acids (FFA), products of lipolysis, were also examined. We did not observe any significant differences in glycerol, FFA, or serum TG in STAT5^AKO^ female mice (Figure 4). Similar observations were made in male STAT5^AKO^ mice (Figures S5A-C), We also performed *ex-vivo* lipolysis assays on iWAT from STAT5^AKO^ mice and did *not* observe any alterations in basal or adrenergic-induced lipolysis between the two genotypes (Figure 4D). Male STAT5^AKO^ mice had a modest but not statistically significant decrease in adrenergic-stimulated lipolysis (Figure S5D). ^14^C-glucose incorporation into TGs of inguinal adipose tissue was measured ex-vivo (Figure 4E). We did not observe differences in basal or insulin-stimulated TG production in STAT5^AKO^ females as compared to floxed littermate controls. In male mice, we observed greater insulin-stimulated ^14^C-glucose incorporation into TGs of inguinal adipose tissue of floxed controls that was reduced in STAT5^AKO^ mice (Figure S5E). Overall, these data indicate sex-specific differences in lipid metabolism in STAT5^AKO^ mice.

**Figure 4.**
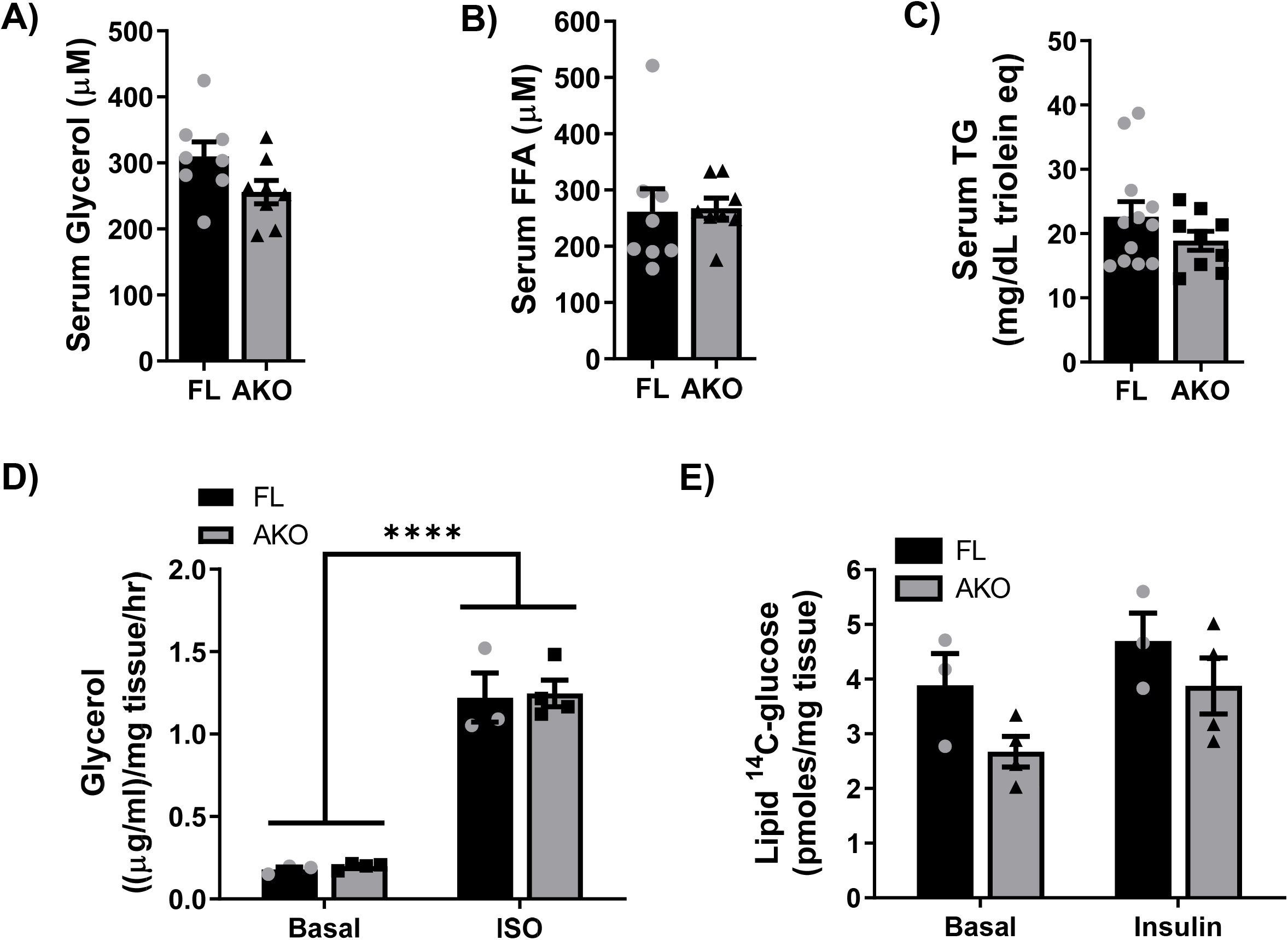
Female STAT5^AKO^ mice do not show any differences in adipose tissue lipid metabolism relative to floxed control mice. Female STAT5^AKO^ (AKO) and floxed (FL) littermate control mice were fed a defined-composition low fat diet (LFD - 10% kcal from fat; A and B) or regular chow (13% kcal from fat; C – E). A and B) After 1 month on LFD diet and at 3 months of age, serum glycerol and free fatty acid (FFA) levels were measured in blood collected from 4 h-fasted mice. C) Blood was collected from 8-week-old, overnight-fasted mice and serum triglyceride (TG) levels were measured. *Ex vivo* lipolysis (D) and *de novo* lipogenesis (E) assays were performed using gWAT and iWAT explants, respectively, from chow-fed mice (5 months old). D) Glycerol release from gWAT explants (~20mg) into media following a 2 h-incubation period was measured under both basal and isoproterenol (ISO)-stimulated (10μM) conditions. E) Incorporation of ^14^C-glucose into total triglycerides was measured by incubating iWAT explants (~50mg) with 4μCi/ml of [^14^C]-U-glucose for 4.5 hours. The triglyceride (neutral lipid) fraction was purified and [^14^C] counts were measured by scintillation counting. For A – C, *t*-tests were used to test for significant between means (n = 8-12 mice per genotype). D and E) Two-way ANOVA with Tukey’s post-hoc multiple comparison analysis was used to test for significance between genotypes and treatments (n = 3-4 mice per group). **** denotes p < 0.0001. See also Figure S4 and Figure S5.

### Sex-specific difference in energy expenditure and adipose tissue gene expression in STAT5^AKO^ mice

Obesity can be associated with alterations in energy expenditure. We singly housed mice for four days prior to transfer to metabolic cages (Promethion Metabolic Analyzer, Sable Systems) to acclimate them to conditions that mimicked our energy expenditure hardware. Mice were placed in fully operational metabolic cages for two days to adapt to new surroundings and then energy expenditure was assessed for three days. As shown in Figure 5A, female STAT5^AKO^ mice had reduced daily energy expenditure (STAT5^AKO^, 5.88 ± 1.42 and FL, 6.75 ± 0.95 kcal/day, p=0.03) when corrected by ANCOVA for lean body mass. As we have observed in all our cohorts of STAT5^AKO^ mice, there were no changes in food intake (Figure 5D). There was increased activity in the dark cycle, but no differences in activity between STAT5^AKO^ mice and floxed littermate controls (Figure 5E). In a separate experiment male STAT5^AKO^ and floxed littermate control mice were placed in metabolic cages for 7 days and provided a standard rodent chow diet *ad libitum*. Figures S6A and S6B demonstrate no differences between groups (STAT5^AKO^, 9.72 ± 0.72 & FL, 9.74 ± 0.70 kcal/day, p=0.40) in total energy expenditure during the experiment. A comparison of substrate utilization (Figure S6C) by respiratory exchange ratio (RER) revealed a significant increase in STAT5^AKO^ males during the light cycle (p=0.01). We calculated energy balance (Figure S6D) by subtracting total daily energy expenditure by energy consumed (kcal). Energy balance was slightly, but significantly increased in STAT5^AKO^ male mice (−1.13 ± 0.9 kcal) versus littermate controls (−1.51 ± 1.0, p= 0.001).

**Figure 5.**
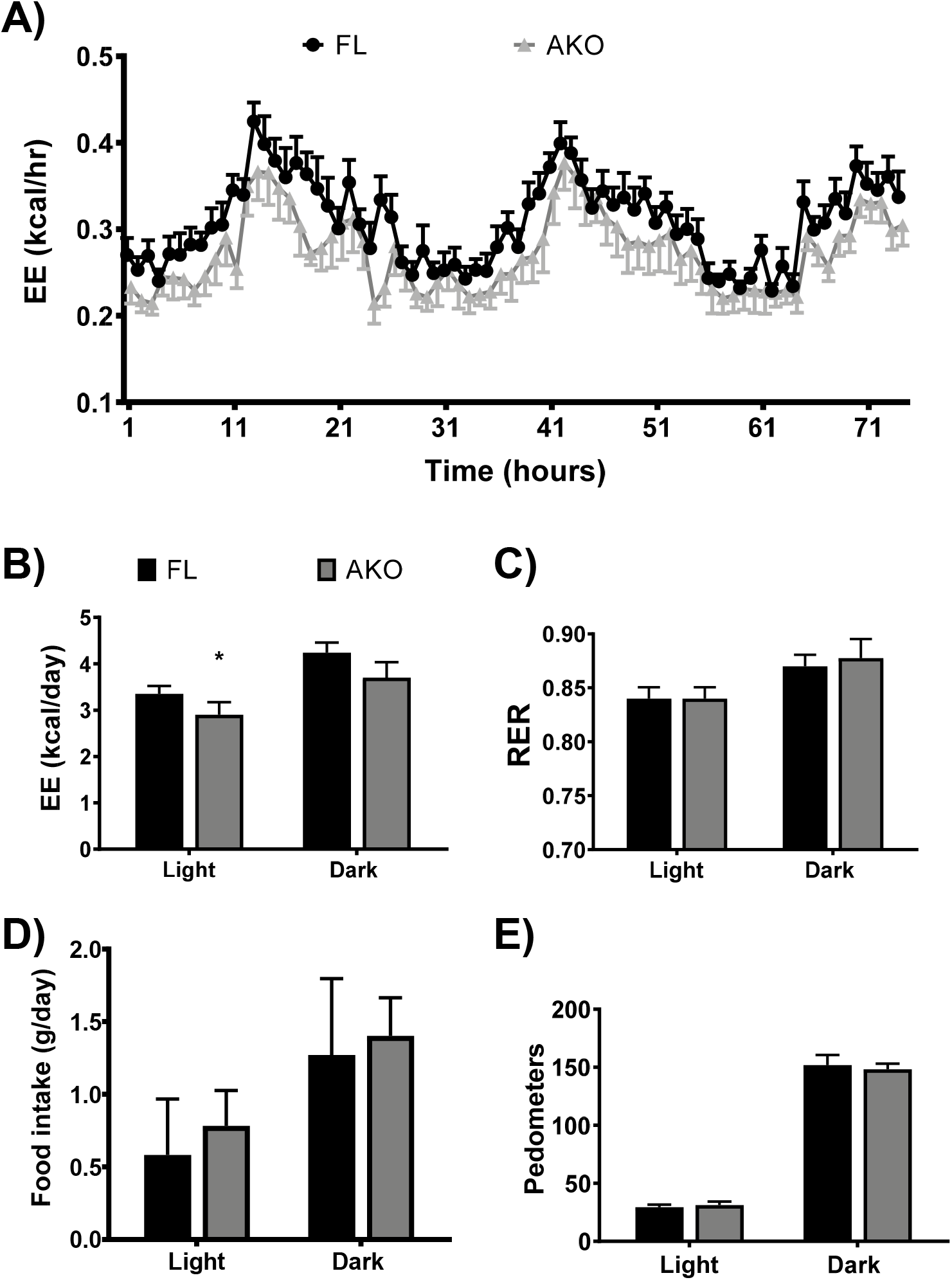
Female STAT5^AKO^ mice have reduced energy expenditure on low fat diet without differences in energy intake or activity. A-C) Total energy expenditure (EE) and respiratory exchange ratio (RER) for floxed (FL) and STAT5AKO (AKO) mice on a low-fat diet (LFD) measured for 3 days. D and E) Food intake and total activity in walking meters (pedometers) was measured during the light and dark cycle. * denotes p < 0.05 (n = 8/group). See also Figure S6.

To investigate molecular mechanisms that may be regulated by loss of STAT5 transcriptional activity in inguinal fat, we performed mRNA sequencing on whole iWAT. The expression changes resulting from loss of adipocyte STAT5 were poorly correlated between females and males (R^2^ = 0.17) (Figure 6A). Furthermore, at each threshold of nominal p-value from 0.05 to 5E-6, the number of differentially regulated genes was consistently higher for males than for females (Figure 6B). Based on an adjusted p-value < 0.05, we identified 387 differentially regulated transcripts in males (289 upregulated and 98 downregulated), whereas only 42 differentially regulated transcripts were identified in females at the same significance threshold (37 upregulated and 5 downregulated). However, the majority of the female differentially expressed transcripts were also differentially expressed in males, resulting in a statistically significant overlap (Fisher exact test p < 2.2E-16) between the two lists (Figure 6C). Thus, despite the broad differences in gene regulation between the two sexes, loss of adipocyte STAT5 also produced some concordant, sex-independent changes in gene expression.

**Figure 6.**
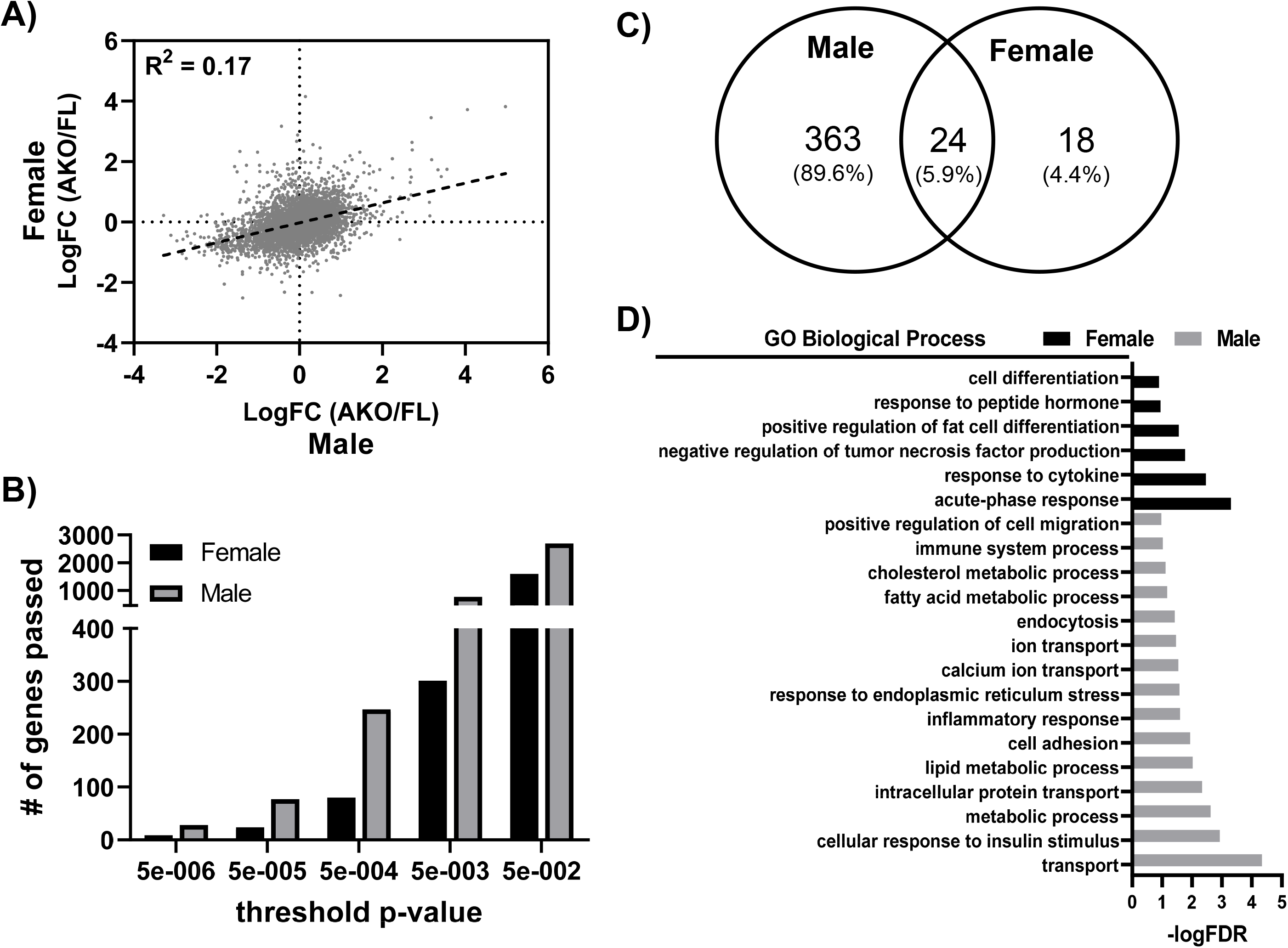
Loss of adipocyte STAT5 results in sexually dimorphic gene expression changes within subcutaneous WAT. Inguinal WAT was collected from male and female STAT5AKO and floxed (FL) control mice at 6 weeks of age, and isolated mRNA was subjected to RNA-sequencing analysis. A) Scatterplot showing correlation between female and male gene expression changes resulting from adipocyte-specific loss of STAT5 (LogFC = Log fold change). The black dashed line is the trend line, and the correlation coefficient (R2) is shown. B) Histogram showing the number of genes differentially regulated at each threshold p-value. C) Venn diagram of differentially regulated genes in male and female mice (AKO/FL) at threshold adjusted p-value equal to 0.05. The percentages of 405 total genes differentially regulated based on p-value criteria are shown for those in common and unique to males or females. D) Genes with an adjusted p value < 0.1 for males and 0.2 for females when comparing AKO versus floxed mice were subjected to DAVID gene ontology (GO) pathway analysis using the following criteria: pathway consists of at least 10 genes for males or 5 genes for females and EASE (expression analysis systematic explorer) score < 0.01. Significance of pathway enrichment is plotted on the x-axis as the negative logarithm of the false discovery rate (FDR).

Gene ontology (GO) based pathway enrichment analysis demonstrated that the most affected biological processes in females were related to cell differentiation, and response to peptide hormones or cytokines (Figure 6D). A notable limitation of this RNA sequencing analysis was that it was performed on entire the iWAT depot where over half of the cells in fat are not adipocytes (Lee et al., 2013; Roh et al., 2017) and as demonstrated in Figure 1F a compensatory increase in SVF STAT5 levels was observed. Although female STAT5^AKO^ mice demonstrate the most differential gene expression in processes more related to non-adipocyte cells, the sexual dimorphism is intriguing. Male mice also have some biological processes that may be altered due to gene expression differences in SVF cells (e.g. positive regulation of cell migration, immune system process, inflammatory response, and cell adhesion); however, the loss of adipocyte STAT5 did not show any common findings for significantly enriched biological processes between females and males (adjusted p <0.1 level). Notably, GO analysis for male STAT5^AKO^ mice revealed biological processes important in adipocyte function and lipid metabolism, such as cellular response to insulin, metabolic process, lipid metabolic process, fatty acid metabolic process, and cholesterol metabolic process. Overall, these RNA sequencing data revealed largely sex-dependent gene expression changes in whole iWAT resulting from loss of adipocyte STAT5, with a greater transcriptomic response observed in males.

### Adipocyte STAT5 is critical for GH-induced insulin resistance, but disposable for GH-induced fat mass loss

Growth hormone is well known for its ability to reduce adipose tissue mass and to promote insulin resistance when administered chronically. Although STAT5^AKO^ mice on chow or low-fat diets have increased subcutaneous adiposity, these differences in adiposity are lost with high fat or high fat-high sucrose feeding (data not shown). After high fat/high sucrose feeding, male STAT5^AKO^ mice and littermate controls had equivalent body mass and fat mass. Mice were injected daily with either saline vehicle or mGH (1.5 mg/kg) subcutaneously for 30-40 days while maintained on regular chow diet. As shown in Figure 7 (A and B), floxed mice with chronic GH treatment were the most insulin resistant as determined by an insulin tolerance test. Saline-treated floxed mice were more insulin sensitive than GH-treated floxed mice. However, STAT5^AKO^ mice were the most insulin sensitive and there was no effect of GH on insulin-induced glucose clearance in these mice (Figures 7A and 7B). We also monitored body weight and fat mass over the 40-day experimental period. Although all mice lost some fat mass because of handling and daily injections, chronic GH treatment, as expected, significantly reduced adiposity when compared to saline-treated mice (Figures 7C and 7D). Surprisingly, GH-treated STAT5^AKO^ mice demonstrated similar fat mass reduction relative to the floxed control mice. Therefore, these data suggest that loss of STAT5 in adipocytes prevents GH-induced systemic insulin resistance and that GH mediates fat loss in a manner independent of adipocyte STAT5.

**Figure 7.**
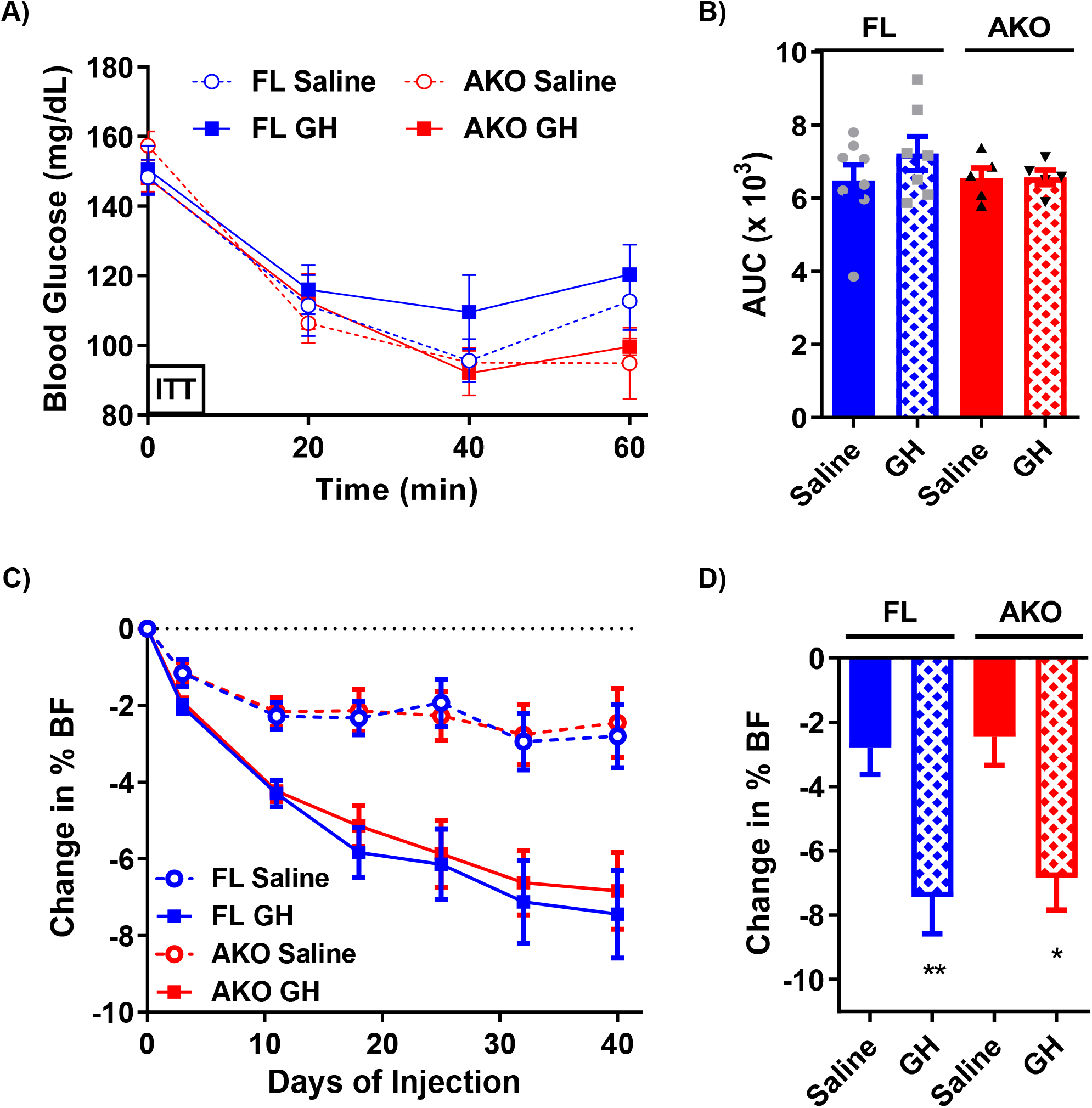
Adipocyte STAT5 is necessary for the ability of GH to induce insulin resistance, but not for its ability to induce loss of body fat. Male STAT5^AKO^ (AKO) and floxed (FL) control mice were fed high fat high sucrose (HFHS; 45% kcal from fat, 30% from sucrose) for 8 weeks beginning at 7 weeks of age, followed by breeder chow (26% kcal from fat) for 7 weeks, and then regular chow diet (13% kcal from fat) for 16 weeks. Following this diet regimen, at 33 weeks of age (and 11 weeks on regular chow), the mice were treated with 0.9% saline or mGH (1.5 mg/kg) for 30-40 days while maintained on regular chow diet. A) Insulin tolerance tests (ITTs) were performed on day 30 of chronic saline/GH injections. Each mouse was injected with 1 U/kg insulin intraperitoneally and blood glucose values were measured at the indicated times. B) Area under the curve (AUC) values were calculated from the ITT curves shown in A. C) The change in percent body fat (BF) or adiposity relative to day 0 was measured over the course of chronic injections, and the total change in % body fat on day 40 is shown in D. Significance was determined by t-test and is denoted as * p < 0.05 and ** p< 0.01 for comparisons between saline and GH treatments (n = 5 – 8 per group).

## DISCUSSION

STAT5 is known to be an important modulator of adipocyte development [reviewed in (Richard and Stephens, 2014)], but the cell-specific contributions of STAT5 are still poorly understood. As STAT5 is a primary mediator of GH signaling, it is important to understand the biological contributions of STAT5 to fat cell functions *in vivo* and how it impacts systemic metabolic health. To investigate the physiological roles of STAT5 in adipocytes, we generated mice lacking both STAT5 genes in adiponectin-expressing cells (STAT5^AKO^). Another group has also generated this mouse model and in male mice, they observed increased adiposity, increased adipocyte size, and evidence of improved insulin sensitivity (Kaltenecker et al., 2017). An advantage to our study is that we examined both male and female mice. We show that male and female STAT5^AKO^ mice have a prominent subcutaneous adiposity phenotype on chow diets and have indicators of improved insulin sensitivity (Figures 2 and 2S). Notably, other interesting observations of STAT5^AKO^ mice were not predicted and provide additional insight into GH action on adipocytes that challenge the current dogma. These unpredicted phenotypic observations in the STAT5^AKO^ mice include: 1) the increased adiposity is not consistent with changes in lipolysis; 2) exogenous murine GH administration reduces fat mass similarly in STAT5^AKO^ and floxed control mice; and 3) significant sex-specific differences in energy expenditure and gene expression are observed in STAT5^AKO^ mice. These observations reveal fundamental gaps in our understanding of adipocyte STAT5 and its ability to contribute to systemic metabolic regulation.

Obesity is associated with altered hypertrophy and/or hyperplasia. As shown in Figures 3 and S3, hypertrophy of both iWAT and gWAT occurs in female and male STAT5^AKO^ mice. Although there are no changes in gWAT mass in STAT5^AKO^ mice (Figures 2 and S2), we observed a substantial increase in adipocyte size in both depots that was accompanied by a decrease in the number of smaller adipocytes (Figures 3C, S3C, 3F and S3F). These data suggest that the lack of GH signaling through STAT5 remodels gonadal fat without affecting depot mass and shows depot-specific effects that occur in both males and females.

In mice that congenitally lack GH, GH has a more pronounced effect on subcutaneous WAT (List et al., 2019a). These observations are consistent with our observations of increased inguinal, but not gonadal, fat mass. Mice that lack GHR in adipocytes, have increased adipocyte size and increased WAT mass (List et al., 2019b), similar to our observations in STAT5^AKO^. Of note, mice that lack GHR in adipocytes also have increased adipocyte size in most depots, but not in male gonadal fat. This is different than adipocyte-specific STAT5 knockout mice, which have increased adipocyte size in inguinal and gonadal fat in males and females. The increased adipocyte size in gonadal fat is not accompanied by increased fat mass (Figures 2 and S2), unlike inguinal subcutaneous fat which is significantly increased. These data indicate that GH is an important modulator of energy metabolism independent of growth effects that are mostly mediated by IGF-1.

The profound effects of GH on glucose metabolism have been known for over a century (Ranke and Wit, 2018) and its diabetogenic effects were first observed in 1930 (Houssay and Biasotti, 1930). GH regulates several metabolic pathways including lipolysis, lipogenesis, glucose uptake, and protein synthesis (Møller et al., 1995). Mice with GH excess are lean and resistant to diet-induced obesity but are severely insulin-resistant (Basu et al., 2018; Berryman et al., 2004; Olsson et al., 2005). In contrast, mice with chronic GH deficiency have increased insulin sensitivity despite increased adiposity (Berryman et al., 2004; Coschigano et al., 2003; Dominici et al., 2002). Increased adipose tissue mass is present in GH deficiency in children and adults, and treatment of GH-deficient patients with exogenous GH can improve lean body mass while reducing fat mass (Ranke and Wit, 2018). The ability of GH to reduce adipose tissue mass had been primarily attributed to its lipolytic effects. Male mice that lack growth hormone receptor (GHR) (List et al., 2013), JAK2 (Corbit et al., 2017b, 2018, 2019; Nordstrom et al., 2013) or STAT5 (Kaltenecker et al., 2017) specifically in adipocytes have been previously studied. Each of these mouse models have increased adiposity that is attributed to decreased lipolytic rates. In terms of potential mechanisms, impaired lipolysis in STAT5^AKO^ mice has been attributed to decreased ATGL and CGI-58 levels in subcutaneous fat (Kaltenecker et al., 2017).

Our findings contrast with these published data. Our STAT5^AKO^ mice do not have significant changes in circulating levels of glycerol or NEFAs, the products of lipolysis (Figures 4 and S5). While our STAT5^AKO^ male mice do have significantly decreased mRNA expression of lipolysis-related genes (ATGL, CGI-58, and ADRB3), only CGI-58 protein expression is significantly decreased in male and female STAT5^AKO^ mice (Figure S4). However, in our model reduced CGI-58 protein levels do not correlate with diminished adipose tissue lipolysis in either sex as we do not observe differences between floxed and STAT5^AKO^ mice in circulating glycerol and NEFA levels or basal or adrenergic-stimulated lipolysis in *ex-vivo* studies (Figures 4 and S5B).

Another consideration is that GH promotes beige fat formation via GHR-induced STAT activation (Nelson et al., 2018). Although altered cold tolerance has been reported in STAT5^AKO^ mice (Kaltenecker et al., 2020), our STAT5^AKO^ mice did not exhibit any differences in acute cold tolerance over an 8-hour period (data not shown). Of note, the iWAT in our STAT5^AKO^ is whiter in appearance in both male and female STAT5^AKO^ (Figures 2 and S2), suggesting there might be decreased beiging gene expression. However, we did not observe alterations in UCP-1 expression or other beige markers (data not shown) in iWAT which is consistent with our data indicating no alterations in cold tolerance. Although BAT demonstrates the most substantial decrease in STAT5 expression (Figures 1C, 1D, and S1C), both male and female mice maintain increased adiposity under thermoneutral conditions (Figures 2H and S2H) suggesting that the increased fat mass of STAT5^AKO^ mice is not due to altered BAT thermogenesis.

These data bring up at least two important questions. First, why are our observations divergent from those in the other STAT5^AKO^ mice? Second, what cellular pathway(s) confer the increased iWAT mass in mice lacking STAT5 in adipocytes? There are a variety of factors that could account for the difference in phenotypic observations ranging from the genetic drift in mouse strains, animal housing conditions, as well as potential contributions of gut microbiota. Also, our fasting conditions are limited to 4 hours or overnight for some studies, while the observations of increased lipolytic products in serum of adipocyte GHR, JAK2, or STAT5 KOs occur after prolonged fasts ranging up to 48 hours (Corbit et al., 2017b, 2018, 2019; Kaltenecker et al., 2017; List et al., 2013; Nordstrom et al., 2013). It should also be noted that mice lacking the lipolytic mediator, ATGL, specifically in adipocytes have a very different phenotype than mice lacking STAT5 in adipocytes. This observation supports our data and suggests that STAT5^AKO^ mice are not obese due to diminished adipocyte lipolysis.

Although it is not evident which lipid metabolism pathway accounts for the increased fat mass in STAT5^AKO^ mice on chow diet, it was very surprising to observe that the increased adiposity of STAT5^AKO^ mice is lost with high fat feeding or high fat/high sucrose feeding (data not shown) and that chronic GH administration reduces fat mass in the same manner in STAT5^AKO^ males as compared to floxed littermate controls (Figures 7C and 7D). These data indicate that the contributions of STAT5 to adipose tissue expansion and loss are clearly different. Our data also suggest that in females, the increased adiposity is likely due, at least in part, to the decreased energy expenditure as we do not observe differences in food intake in STAT5^AKO^ mice.

Acromegaly is a hormonal disorder in which GH levels are chronically elevated and cannot be suppressed by typical feedback mechanisms. Its primary metabolic consequence is insulin resistance (Vila et al., 2019), which is also observed in transgenic mouse models of GH excess (Benencia et al., 2015; Berryman et al., 2004; Olsson et al., 2005). GH-induced insulin resistance has been attributed to the pro-lipolytic action of GH because increased FFA levels are associated with decreased insulin sensitivity in other physiological and pathophysiological metabolic contexts (Karpe et al., 2011; Nielsen et al., 2001; Vila et al., 2019). The strongest and most direct evidence for this comes from a small study in a group of 7 adult-onset GH-deficient patients who were given four separate treatments of GH and/or the antilipolytic agent acipimox at different times followed by assessment of insulin resistance via hyperinsulinemic/euglycemic clamp (Nielsen et al., 2001). Acipimox administration substantially attenuated insulin resistance in GH-treated patients demonstrating a causal link between GH-induced lipolysis and insulin resistance; however, the authors made note that GH was still able to slightly decrease insulin sensitivity in the presence of acipimox suggesting that GH may also promote insulin resistance through non-lipolytic/non-FFA-dependent mechanisms (Nielsen et al., 2001). Interestingly, these findings are supported by our data from STAT5^AKO^ mice following chronic GH treatment. The results in Figure 7 demonstrate that the effect of GH on insulin resistance and lipolysis-induced fat mass reduction can be dissociated by the loss of STAT5 in adipocytes and thereby provide evidence of a potential non-FFA-dependent molecular mechanism, mediated by adipocyte STAT5, by which GH can reduce insulin sensitivity in conditions of GH excess, such as acromegaly.

A major challenge with the GH literature is interpreting studies that use bovine GH transgenes (Benencia et al., 2015; Berryman et al., 2004; Householder et al., 2018; Olsson et al., 2005) in mice or use bovine or human GH (Ding et al., 2011; Kim et al., 2012; List et al., 2009) in mice. It is not apparent that these responses are physiological or due to GH receptor signaling. Unlike insulin and other hormones, GH does not widely cross species boundaries (Goodman et al., 1996). Therefore, to study actions of GH in mice, we used murine GH (mGH). We confirmed that mGH acutely activates GHR signaling and induction of a STAT5 target gene (Figure 1E). After confirming the bioactivity of mGH, we examined the ability of mGH to induce insulin resistance in obese male mice that had been exposed to high-fat diets. Following high-fat feeding, STAT5^AKO^ mice do not have a difference in adiposity like they do on chow diets (data not shown). After high fat/high sucrose feeding, both genotypes had equivalent body weights and fat mass at the start of the chronic GH treatment experiment shown in Figure 7. These studies revealed that chronic GH treatment induced insulin resistance, as assessed by ITT, in floxed littermates, but not in STAT5^AKO^ mice. STAT5^AKO^ mice are insulin-sensitive, and they are protected from GH-induced insulin resistance. These data suggest that GH-induced insulin resistance is mediated, at least in part, by adipocyte STAT5 expression.

In summary, investigation of STAT5^AKO^ mice revealed several intriguing observations. Despite the gender-independent changes in subcutaneous fat mass expansion, HOMA-IR, as well as increased adipocyte size in subcutaneous and visceral depots, STAT5^AKO^ mice exhibit prominent sex-specific differences. The most prominent similarity of STAT5^AKO^ male and female mice is their increased adipose tissue mass that is maintained at thermoneutrality (Figures 2 and S2), indicating the robustness of the adiposity phenotype in STAT5^AKO^ mice. The increased fat mass in female STAT5^AKO^ mice is associated with decreased energy expenditure (Figure 5), but not in changes in lipolytic gene expression (Figure S4) or changes in lipolysis (Figure 4). In male STAT5^AKO^ mice, we observed greater changes in gene expression in whole inguinal adipose tissue (Figure 6), but no changes in energy expenditure (Figure S6). Divergences in both energy expenditure and gene expression are prominent differences and suggest that male and female STAT5^AKO^ mice develop increased adiposity as the result of different sex-dependent mechanisms. Based on other animal models with disrupted adipose tissue GH or STAT5 signaling (Kaltenecker et al., 2017; List et al., 2013, 2019b; Ran et al., 2019), we expected GH-induced weight loss to be dependent on adipocyte STAT5, but STAT5^AKO^ mice have an equivalent reduction in AT mass (Figure 7). Hence, in this model, GH-induced loss of fat mass is not dependent on STAT5 even though there is clear evidence that adipocyte STAT5 contributes to AT deposition. Other known mediators of GH action, such as the phospholipase C/protein kinase C (PLC/PKC) and MEK/ERK are potential candidates as these pathways have been previously shown to contribute to GH-induced lipolysis in rainbow trout hepatocytes (Bergan et al., 2013) and murine (Sharma et al., 2018a) and human (Sharma et al., 2018b) adipocytes. In fact, it was recently shown that GH can stimulate lipolysis by activating the MEK/ERK pathway and inhibiting PPARγ signaling, which results in decreased expression of a negative regulator of lipolysis, Fat Specific Protein 27 (FSP27) (Sharma et al., 2018a, 2018b). Interestingly, this study demonstrated a complex relationship between STAT5 activation by GH, FSP27 expression, and lipolysis. Pharmacological inhibition of STAT5 phospho-activation in the presence of GH reduced FSP27 expression and increased lipolysis more than observed with GH alone (Sharma et al., 2018a) suggesting that STAT5 may actually play an antilipolytic role in the context of GH signaling. While we observed increased ERK phosphorylation in STAT5^AKO^ male and female mice (Figure S4B and S4C), *Fsp27* (*Cidec*) expression was unaltered in our RNA-seq analyses (GEO: GSE113939). Clearly, these observations merit further study to determine which GH signaling pathway(s) account for STAT5-independent weight loss in a model of GH excess and to understand the complex nature of the crosstalk between signaling pathways that ultimately give rise to increased adiposity in the absence of adipocyte STAT5 that is distinct in males and females. In addition to our unexpected observations, these studies indicate that STAT5^AKO^ mice are a useful tool to investigate sexspecific differences in adiposity, GH signaling, and adipose tissue depot-specific responses.

## Supporting information

Key Resources Table

## ACKNOWLEDGMENTS

We extend our sincere appreciation to Tamra Mendoza for providing support for all our animal experiments. We are also grateful to the Pennington Biomedical Research Center Transgenics Core for generating the STAT5^AKO^ mice, and to Dr. David Burk for his assistance with adipocyte size analysis. GH was purchased from the National Hormone & Peptide Program (Scientific Director - Dr. A.F. Parlow). This research project utilized the facilities of Genomics Core, the Cell Biology and Bioimaging Core, and the Animal Metabolism and Behavior Core at Pennington Biomedical that are supported in part by COBRE (1P30GM118430) and NORC (NIH 2P30DK072476) center grants from the National Institutes of Health. The Promethion Metabolic cage system was purchased using funds from NIH shared instrumentation grant S10OD023703. C.M.E. is supported by R03 DK122121 from NIH. This work was supported by NIH grant R01DK052968 (J.M.S.) and pilot funding to C.M.E. from a NORC center grant P30DK072476.

## AUTHOR CONTRIBUTIONS

Conceptualization, A.J.R., C.M.E. and J.M.S.; Methodology, A.J.R., C.M.E., H.H., and P.Z.; Investigation, A.J.R., C.M.E., H.H., and P.Z., Formal Analysis, A.J.R., H.H., T.D.A., and S.G.; Writing – Original Draft, J.M.S. and A.J.R.; Writing – Review & Editing, all authors; Visualization, A.J.R., H.H., T.D.A., and S.G.; Supervision, J.M.S., A.J.R., and C.M.E.; Funding Acquisition, J.M.S. and C.M.E.

## STAR ★ METHODS

Detailed methods are provided in the online version of this manuscript and include the following:

- KEY RESOURCES TABLE
- CONTACT FOR REAGENT AND RESOURCE SHARING
- EXPERIMENTAL MODEL AND SUBJECT DETAILS
  ○ Animals
  ○ Chronic GH treatment study

- METHODS DETAILS
  ○ Body composition measurements (NMR)
  ○ Blood glucose and serum analyses
  ○ RNA isolation and RT-qPCR quantification
  ○ Immunoblotting
  ○ Adipose Tissue Fractionation
  ○ Adipose histology and adipocyte size analysis
  ○ *Ex vivo* lipolysis
  ○ *Ex vivo de novo* lipogenesis
  ○ Indirect calorimetry/energy expenditure analyses
  ○ RNA-sequencing and analysis
  ○ Insulin tolerance test

- QUANTIFICATION AND STATISTICAL ANALYSIS
- MATERIALS AVAILABILITY
- DATA AND SOFTWARE AVAILABILITY

## STAR ★ METHODS

### CONTACT FOR REAGENT AND RESOURCE SHARING

Further information and requests for resources and reagents should be directed to and will be fulfilled by the Lead Contact, Jacqueline Stephens.

### EXPERIMENTAL MODEL AND SUBJECT DETAILS

#### Animals

We obtained floxed *Stat5a/b* (Stat5^fl/fl^) breeding pairs on a C57BL/6 background from Dr. Lothar Hennighausen. In these transgenic mice, a set of loxP sites flanks both *Stat5a* and *Stat5b* genes. We crossed Stat5^fl/fl^ mice with adiponectin-Cre (Adipoq-Cre) mice to generate adipocyte specific STAT5 knockout (STAT5fl/fl; AdipoQ-Cre) mice that we refer to as STAT5^AKO^ mice. Unless otherwise stated, mice were housed in a temperature (22 ± 2 °C)- and humidity-controlled (45–55%) room under a 12-h light/dark cycle with free access to food and water. For experiments at thermoneutrality, mice were housed at 28 °C. For the studies, mice were used between 6 weeks to 11 months of age and fed standard chow (LabDiet®5001; 13% kcal from fat, 57% from carbohydrates, and 30% from protein), breeder chow (LabDiet®5015; 26% kcal from fat, 54% from carbohydrates, and 20% from protein), low fat diet, no sucrose (LFD; Research Diets D12450K; 10% kcal from fat, 70% from carbohydrates, and 20% from protein), and/or high fat diet (HFD; Research Diets D12492; 60% kcal from fat, 20% from carbohydrates, and 20% from protein) as indicated in figure legends. All animal care and use procedures were conducted in accordance with the regulations of the Institutional Animal Care and Use Committee at Pennington Biomedical Research Center (under protocol numbers P863, P977 and P985).

#### Chronic GH treatment study

Male STAT5^AKO^ and floxed control mice were fed high fat high sucrose (HFHS) diet (45% kcal from fat, 30% kcal from sucrose; Research Diets D08112601) for 8 weeks beginning at 7 weeks of age, followed by breeder chow (LabDiet®5015) for 7 weeks, and then standard chow diet (LabDiet®5001) for 16 weeks. Following this diet regimen, at 33 weeks of age (and 11 weeks on regular chow), the mice began receiving daily subcutaneous injections of 0.9% saline or mGH (1.5 mg/kg) for 30-40 days while maintained on regular chow diet. Injections were administered at 3:00p ± 2 hours each day, and they were rotated between right and left sides of the back to minimize skin irritations. Body weight and body composition (measured via NMR) were assessed weekly. Following 30 days of GH treatment, insulin tolerance tests were administered to the mice.

### METHODS DETAILS

#### Body composition measurements (NMR)

Animal body composition was measured by NMR (Bruker Minispec LF110). Body weights were measured concurrent with NMR assessments, and adiposity was calculated as total fat mass divided by total body weight x 100.

#### Blood Glucose and serum analyses

Blood samples were collected following a 4-hour fast and serum was collected using a Microtainer®tube with a serum separator additive (BD). Whole blood glucose levels were measured using a Breeze 2 glucometer (Bayer). Serum GH (Millipore Sigma), IGF-1 (Crystal Chem), and insulin (Crystal Chem) were measured by ELISA according to manufacturer instructions. Serum glycerol (Sigma) and NEFA (BioVision) levels were measured using colorimetric assays. Homeostasis model assessment of insulin resistance (HOMA-IR) was calculated from glucose and insulin concentrations using the following formula: fasting glucose (mg/dl) × fasting insulin (μU/ml)/405 (Frøsig et al., 2013; Matthews et al., 1985) For triglyceride (TG) measurements, blood was collected from overnight-fasted mice using Sarstedt Microvette CB300 capillary blood collection tubes to separate the serum. Serum TG levels were measured using a colorimetric kit (Sigma) adapted for spectrophotometric analysis using a microplate reader (VersaMax, Molecular Devices).

#### RNA isolation and RT-qPCR quantification

Tissues were flash frozen in liquid nitrogen and stored at –80 °C. RNA was extracted by homogenizing the tissue in TRIzol reagent (Thermo Fisher) using a S 25 N – 10 G dispersing probe (IKA Works, Inc.). Chloroform extraction was performed according to the TRIzol reagent manufacturer’s protocol to isolate RNA that was further purified using the RNeasy mini kit (Qiagen) and quantified using a NanoDrop spectrophotometer. cDNAs were synthesized with the High-Capacity cDNA Reverse Transcription (RT) kit (Applied Biosystems). SYBR Green Premix (Takara Bio USA) and primers from IDT (Integrated DNA Technologies) were used to perform quantitative real-time PCR (qPCR) on an Applied Biosystems 7900HT system with SDS 2.4 software (Applied Biosystems) and the following thermal cycling conditions: 2 min at 50°C, 10 min at 95°C, 40 cycles of 15 s at 95°C, and 1 min at 60°C with final dissociation stage: 15s at 95°C, 15 s at 60°C, and 15 s at 95°C. Expression levels were determined using the relative standard curve method with non-POU-domain-containing, octamer-binding protein (*Nono*) or cyclophilin a (*Ppia*) used as reference genes. Primer sequences are listed in Table S1.

#### Immunoblotting

Tissues were homogenized in immunoprecipitation (IP) buffer containing 10 mM Tris (pH 7.4), 150 mM NaCl,1 M ethylene glycol tetraacetic acid (EGTA), 1 mM ethylenediaminetetraacetic acid (EDTA), 1% Triton X-100, 0.5% Igepal CA-630 plus protease and phosphatase inhibitors (1 mM phenylmethylsulfonyl fluoride, 1μM pepstatin, 50 trypsin inhibitory milliunits of aprotinin, 10μM leupeptin, 1 mM 10-phenanthroline, and 0.2 mM sodium orthovanadate, and 100μM sodium fluoride). The concentrations of protease and phosphatase inhibitors were respectively doubled and quadrupled for all tissue homogenates. Lysates were clarified by centrifugation at 17,500 x g for 10 min at 4 °C. Following removal of the floating lipid layer, the supernatant was recovered, and protein concentration was determined by a bicinchoninic acid (BCA) protein assay (Sigma).

Protein extracts (50 μg total protein per lane) were separated on 7.5 or 10% sodium dodecyl sulfate (SDS)-polyacrylamide gels, and then transferred to nitrocellulose membranes as previously described (Richard et al., 2014). Membranes were probed according to standard immunoblotting techniques using primary antibodies to STAT5A, STAT5B, ADPN, ATGL, CGI-58, ADRB3, phospho-ERK1/2, and total ERK1/2 as indicated in the Key Resources Table. Primary antibodies were detected using anti-rabbit or anti-mouse horseradish peroxidase (HRP)- conjugated secondary antibodies (Jackson ImmunoResearch) and visualized using SuperSignal West Pico PLUS detection reagents (Thermo Fisher Scientific). Films were scanned, and band intensities were quantified using Image Studio software (LI-COR Biosciences).

#### Adipose Tissue Fractionation

Fractionation of adipose tissue was performed using a protocol adapted from (Grant et al., 2013). Briefly, adipose tissue was dissected, weighed, and immediately placed on ice in DMEM + 5% heat inactivated FBS until it was minced into small pieces. Minced adipose tissue was incubated with Type 1 collagenase (Worthington Biochemical) at 37 °C in a shaking water bath until fully digested (approximately 1 to 1.5 hours) and then centrifuged at 500 x g for 5 min. The floating layer of adipocytes was transferred to a fresh tube and mixed 1:4 with 5X IP buffer, while the supernatant was discarded, and the pellet was resuspended in ACK lysing buffer for red blood cell lysis. The lysis reaction was neutralized with PBS, and the sample was centrifuged at 500 x g for 5min. The resuspended pellet was filtered by successive passages through cell strainers (pore sizes of 100μm and 40μm) and washed with PBS before a final centrifugation at 10,000 x g to yield the stromal vascular fraction (SVF) pellet, which was subsequently resuspended in 1X IP buffer. Following a single freeze/thaw cycle at −80 °C, the resuspended adipocyte fraction and SVF were each passed through a 20-gauge needle 10 times and then centrifuged at 10,000 x g for 10 min. The aqueous layer between the floating lipid and pellet of the adipocyte fraction and the supernatant of the SVF were used for immunoblotting analyses. All centrifugation steps were performed at 4 °C.

#### Adipose histology and adipocyte size analysis

Adipose tissues were dissected and fixed in 10% neutral buffered formalin for a minimum of 24 hours. The Cell Biology and Bioimaging Core (CBBC) at Pennington Biomedical performed tissue processing, paraffin embedding, cryo-sectioning into 5 μm thick sections, and hematoxylin and eosin (H&E) staining. The stained tissue was imaged using a NanoZoomer digital slide scanner (Hamamatsu Photonics, Hamamatsu, Japan) and visualized with the associated software (NanoZoomer Digtial Pathology; NDP.view 2.7.52). Adipocyte area was calculated from the slide images (>3000 cells per mouse per depot were quantified) using Visiopharm software (Visiopharm A/S, Hørsholm, Denmark).

#### *Ex vivo* lipolysis

*Ex vivo* lipolysis assays were performed as described in (Schweiger et al., 2014). Mice were fasted for 4 hours prior to euthanasia and WAT excision. Briefly, gonadal WAT explants (10 – 20 mg, in triplicate) were incubated in 200 μl DMEM containing 2% fatty acid (FA)-free BSA, with or without 10 μM isoproterenol in 96 well plates at 37 °C in a 5% CO_2_ and 95% humidified incubator for two hours. Glycerol release was quantified from the conditioned medium. Explants were weighed following the 2-hour incubation period, and glycerol values were normalized to tissue weight.

#### *Ex vivo de novo* lipogenesis

*Ex vivo* lipogenesis assays were performed as described in (Guilherme et al., 2017; Pedersen et al., 2015). Mice were fasted for 4 hours prior to euthanasia and WAT excision. Briefly, inguinal WAT explants (~50 mg, in triplicate) were incubated in labeling media containing 2% FA-free BSA, 0.5 mM D-glucose, 0.5 mM sodium acetate, 2 mM sodium pyruvate, 2 mM glutamine, 1% (v/v) Pen-Strep, and 4 μCi/ml ^14^C-U-glucose for 4.5 hours at 37 °C in a 5% CO_2_ and 95% humidified incubator. Tubes for insulin-stimulated conditions also contained 1 μM insulin (Sigma-Aldrich). Lipid extraction was performed as described (Pedersen et al., 2015), and incorporation of ^14^C-glucose into the FA moiety of extracted triglycerides was quantified by liquid scintillation counting.

#### Indirect Calorimetry/Energy Expenditure Analyses

For these studies, female and male mice were weaned onto chow diet at 3 weeks of age (WOA). They were switched to a characterized low-fat diet (Research Diets D12450K) or chow diet (LabDiets 5001) at 7 WOA and remained on this diet throughout the indirect calorimetry assessment in the metabolic cage system (Promethion, Sable Systems International). At 9 WOA, multi-housed female mice were moved to training cages that have food hoppers and water dispensers exactly like the actual metabolic cages for 3 days prior to singly housing the mice within the training cages for an addition 4 days. After a total of one week in the training cages, the singly housed mice were moved to the Promethion assessment cages. Assessment of male mice was performed one week after female mice.

Oxygen consumption (VO2), carbon dioxide production (VCO2), body weight, food intake, and physical activity were continuously monitored for 5 days within the chambers that were maintained with 12-hr light/dark cycles. The mice had ad libitum access to food and water while in the training and chamber cages. Average daily and cumulative data were calculated over the final 3 days in the chamber. VO2 and CO2 were measured every 30 minutes and used to determine the respiratory exchange ratio (RER = VCO2/VO2) and energy expenditure (EE = 3.185+1.232 x RER) x VO2). VO2, VCO2, and energy expenditure values were normalized to total body weight. Locomotor activity (pedometers) was determined by calculating the sum of all detectable motion determined using an infrared photocell beam interruption technique (> 1 cm/s along X-, Y-, or Z-axis) over the continuous monitoring period.

#### RNA-sequencing and analysis

For RNA-sequencing analysis, RNA was isolated from subcutaneous inguinal WAT of 6-week-old mice as described for RT-qPCR quantification. RNA quality was confirmed using a Bioanalyzer RNA 6000 chip (Agilent). Samples were verified to have RIN values of >7. Sequencing libraries were constructed using a Quant-Seq 3’ mRNA-Seq Library Prep kit (Lexogen). Each sample was prepared with a unique sample index. Completed libraries were analyzed on the Bioanalyzer High Sensitivity DNA chip (Agilent) to verify correct library size. All libraries were pooled in equimolar amounts and sequenced on the NextSeq 500 sequencer (Illumina) at 75bp forward read and 6bp forward index read. Primary data analysis was performed using the Lexogen QuantSeq pipeline V1.8.8 on the Bluebee platform for quality control, mapping, and read count tables. The raw and processed data are deposited in the GEO database (accession number GEO: GSE113939).

Raw count matrices of RNA sequencing data were obtained via *the Rsubread* (Liao et al., 2019) package in R, and further processed for gene quantification and identification of differentially expressed genes using the *limma* package (Ritchie et al., 2015). Genes with at least one count per million (CPM) reads in 6 or more samples were retained for further analysis, resulting in 15118 genes. Gene counts were log2 transformed and normalized for sequencing depth via the TMM method (Robinson and Oshlack, 2010). The mean-variance relationship of gene-wise standard deviation to average logCPM gene signal was assessed via the ‘voom’ method (Law et al., 2014) to generate precision weights for each individual observation. The logCPM values and associated precision weights were subsequently utilized to generate empirical Bayes moderated t-statistics estimates for identification of differentially expressed genes. To adjust for multiple testing, genes with an adjusted p-value <0.05 were considered to be differentially expressed. Venn analysis of gene lists were conducted via Venny (https://bioinfogp.cnb.csic.es/tools/venny/index.html).

Pathway enrichment analysis on differentially expressed genes was conducted via DAVID (Huang et al., 2009) using pathways from the gene ontology (GO) Biological Process repository. To account for differences in the number of differentially expressed genes between males and females, genes with an adjusted p-value for differential expression <0.1 were used for DAVID analysis in males, whereas the corresponding adjusted p-value filter was set to <0.2 for females. Similarly, GO pathways with 10 or more genes were considered for analysis of male samples, and pathways with 5 or more genes were considered for female samples. The DAVID-specific EASE score was set to <0.01 for both sexes.

#### Insulin Tolerance Test

A baseline blood glucose measurement was obtained via tail nick (0 min), and mice were then injected i.p. with 1 U/kg insulin (Humulin R, Eli Lilly). Blood glucose measurements were obtained via tail nick at 20-, 40-, and 60-min post-injection. All blood glucose measurements were performed using a Bayer Breeze 2 glucometer.

### QUANTIFICATION AND STATISTICAL ANALYSIS

All data were expressed as the mean ± SEM. We used GraphPad Prism 8.0 (GraphPad Software, Inc.) to perform the statistical analyses. For comparisons between two independent groups, a Student’s t test was used and p < 0.05 was considered statistically significant. For comparisons between three or more groups, two-way ANOVA with Tukey’s multiple comparisons testing was performed. Analysis of covariance (ANCOVA) was used to determine differences between groups in energy expenditure and substrate oxidation using corrections for lean body mass and fat mass. All sample sizes, statistical test methods, and p values are listed in the figure legends.

### MATERIALS AVAILABILITY

STAT5^AKO^ mice generated in this study are available from the Lead Contact with a completed Material Transfer Agreement if the mice are still being actively bred at the time. If the STAT5^AKO^ mice are unavailable at the time of the request, they can be easily re-created by breeding STAT5^fl/fl^ mice that we can provide with permission from Dr. Lothar Hennighausen, and Adipoq-Cre mice (B6.FVB-Tg(Adiopq-cre)1Evdr/J), which are available from Jackson Laboratory (Stock No. 028020). We are glad to share the STAT5^AKO^ or STAT5^fl/fl^ mice with reasonable compensation for shipping.

### DATA AND SOFTWARE AVAILABILITY

The RNA-seq data generated during this study are available at in the GEO repository under the accession number GEO: GSE113939.

## SUPPLEMENTAL INFORMATION

Supplemental information includes 6 figures and 1 table and can be found with this article online.

