## Supplemental Figures S1 to S7 and Table S1 for "Loss of STAT5 in adipocytes increases subcutaneous fat mass via sex-dependent and depot-specific pathways"

**Figure S1. Related to Figure 1.**

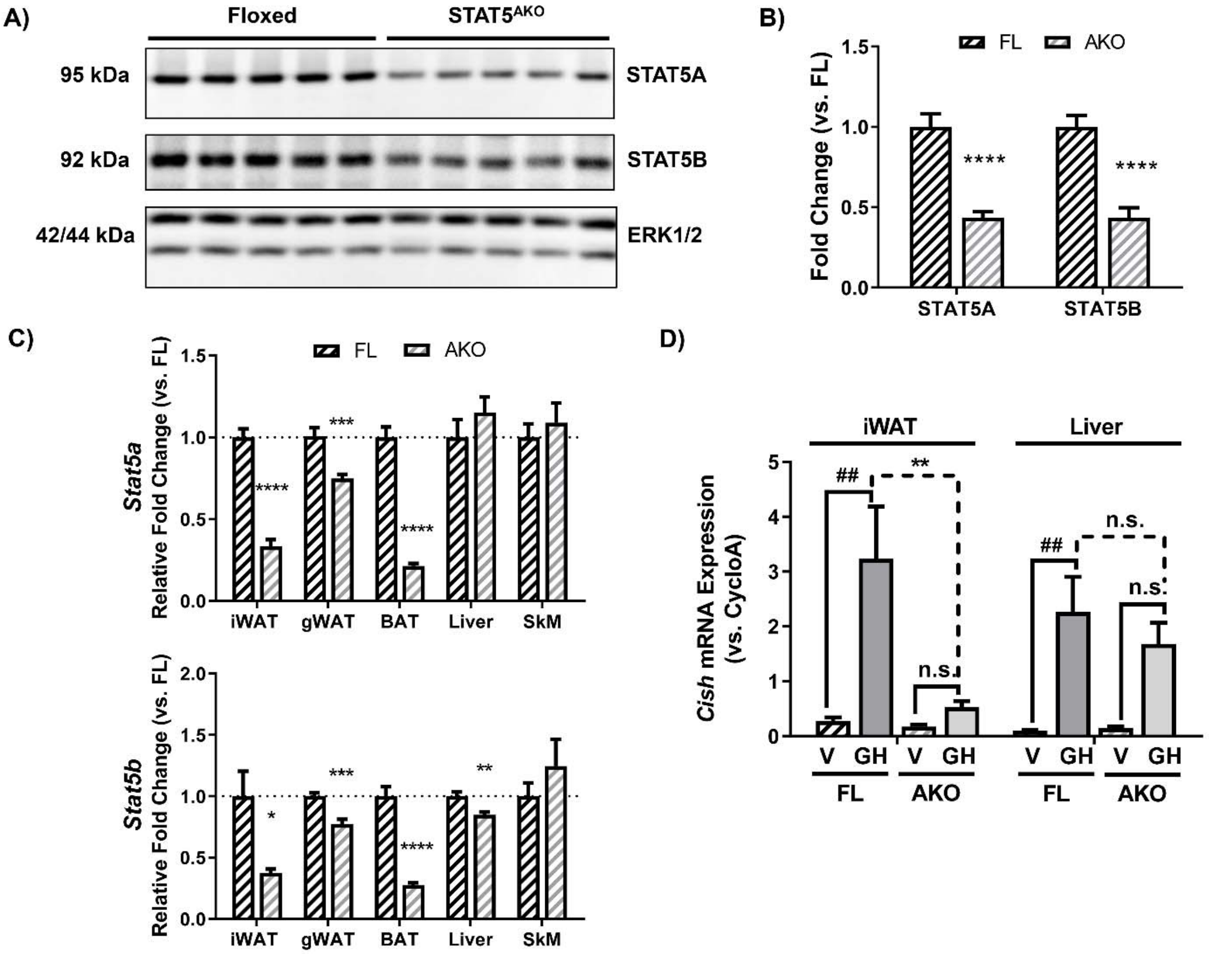


**Figure S1. Expression of STAT5A and 5B is knocked down in adipose tissue and adipocytes of STAT5$^{AKO}$ male mice. Related to Figure 1.** Male STAT5$^{AKO}$ (AKO) mice and their floxed (FL) littermate controls were euthanized at 2 – 3 months of age and tissues were immediately collected for protein or gene expression analyses. A) Immunoblot of proteins resolved from iWAT samples (n = 7 per genotype; 5 representative samples shown for each genotype). B) Quantification of band intensities from A (n = 7 per group). ERK1/2 was used as a loading control, and band intensities were normalized to ERK1/2 and then represented as fold change relative to FL mice. C) *Stat5a (top)* and *Stat5b (bottom)* gene expression measured by RT-qPCR for the indicated tissues (n = 5 – 8 mice per group). D) Mice were injected with 1.5mg/kg mGH or vehicle (V; 0.9% Saline-NaOH) for 30 minutes prior to euthanasia and tissue collection. Gene expression measured by RT-qPCR is shown (n = 5 – 6 mice per group). Significance was determined by *t*-test for AKO versus FL comparisons and is denoted as * $p < 0.05$, ** $p < 0.01$, *** $p < 0.001$, **** $p < 0.0001$, n.s., not significant. For D, a 2-way ANOVA was used to assess treatment/genotype and tissue variables with Tukey's post-hoc multiple comparisons test to compare all treatment/genotype groups for each tissue; $^{\#\#}$ denotes $p < 0.01$ for GH versus V comparisons.

**Figure S2. Related to Figure 2.**

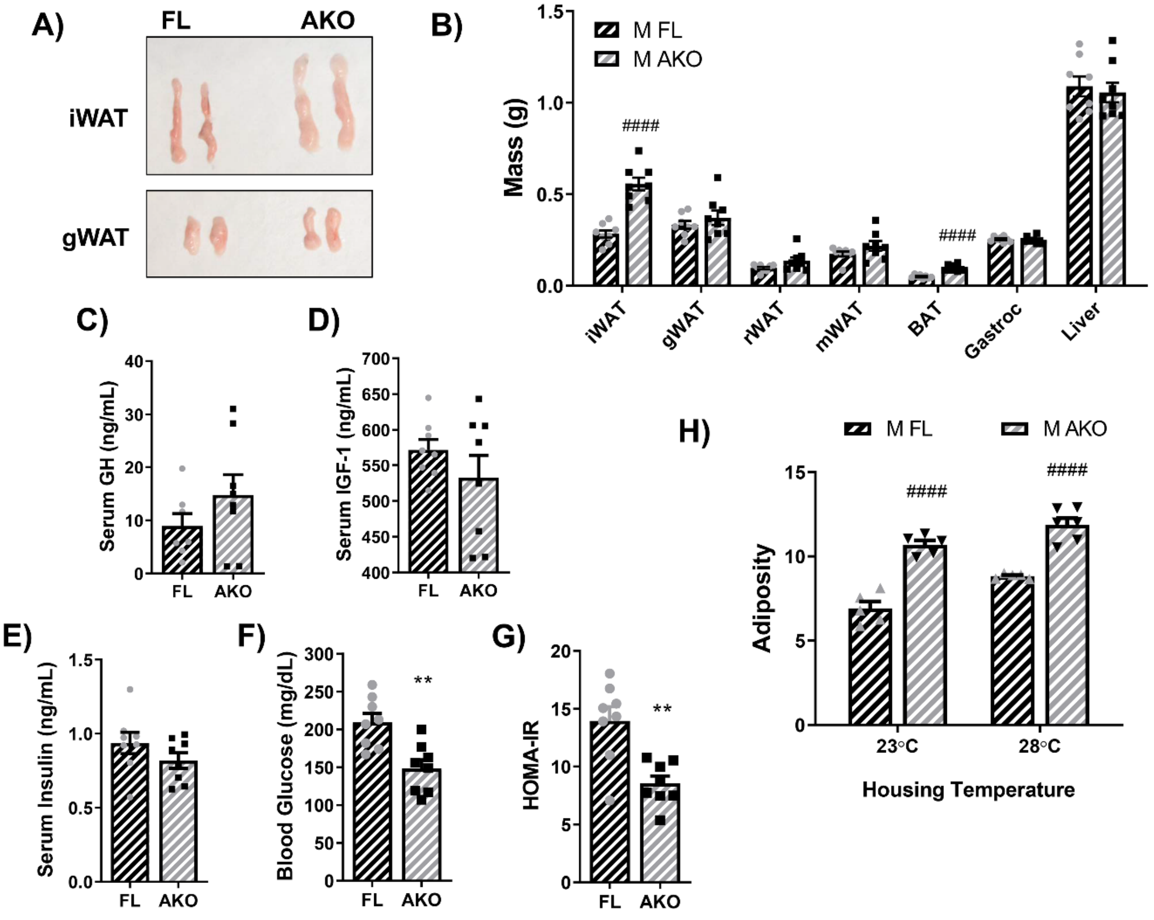


**Figure S2. Male STAT5$^{AKO}$ mice have increased adiposity and are metabolically healthier than floxed controls when fed chow or low-fat diet. Related to Figure 2.** Male (M) STAT5$^{AKO}$ (AKO) and floxed (FL) littermate control mice were weaned onto regular chow diet (13% kcal from fat) and maintained on that diet (A, B, and H) or switched to a defined-composition low fat diet (LFD - 10% kcal from fat; C - G) at 6 weeks of age. A) Representative images of inguinal and gonadal white adipose tissue depots (iWAT and gWAT) collected from five-month-old mice. B) Tissue weights of white adipose tissue depots (iWAT, gWAT, retroperitoneal - rWAT, mesenteric - mWAT), brown adipose tissue (BAT), gastrocnemius skeletal muscle (Gastroc), and liver collected from 10-week-old mice (n = 11-13). C – F) Serum growth hormone (GH), insulin growth factor 1 (IGF-1), insulin, and blood glucose levels collected from 3-month-old mice on LFD for 1 month (n = 7 – 8). G) HOMA-IR was calculated from insulin and glucose levels in E and F. H) Mice (n = 5 – 6 per group) were housed at different temperatures beginning at weaning (3 weeks of age). They were fed chow diet (13% kcal from fat) for 6 weeks and then body weight and body composition (fat mass and lean mass) were measured at 9 weeks of age. Adiposity was calculated as fat mass divided by total body weight for each animal. Significance was determined by *t*-test and is denoted as # $p < 0.05$, ## $p < 0.01$, ### $p < 0.001$, or #### $p < 0.0001$ for AKO versus FL comparisons.

**Figure S3. Related to Figure 3.**

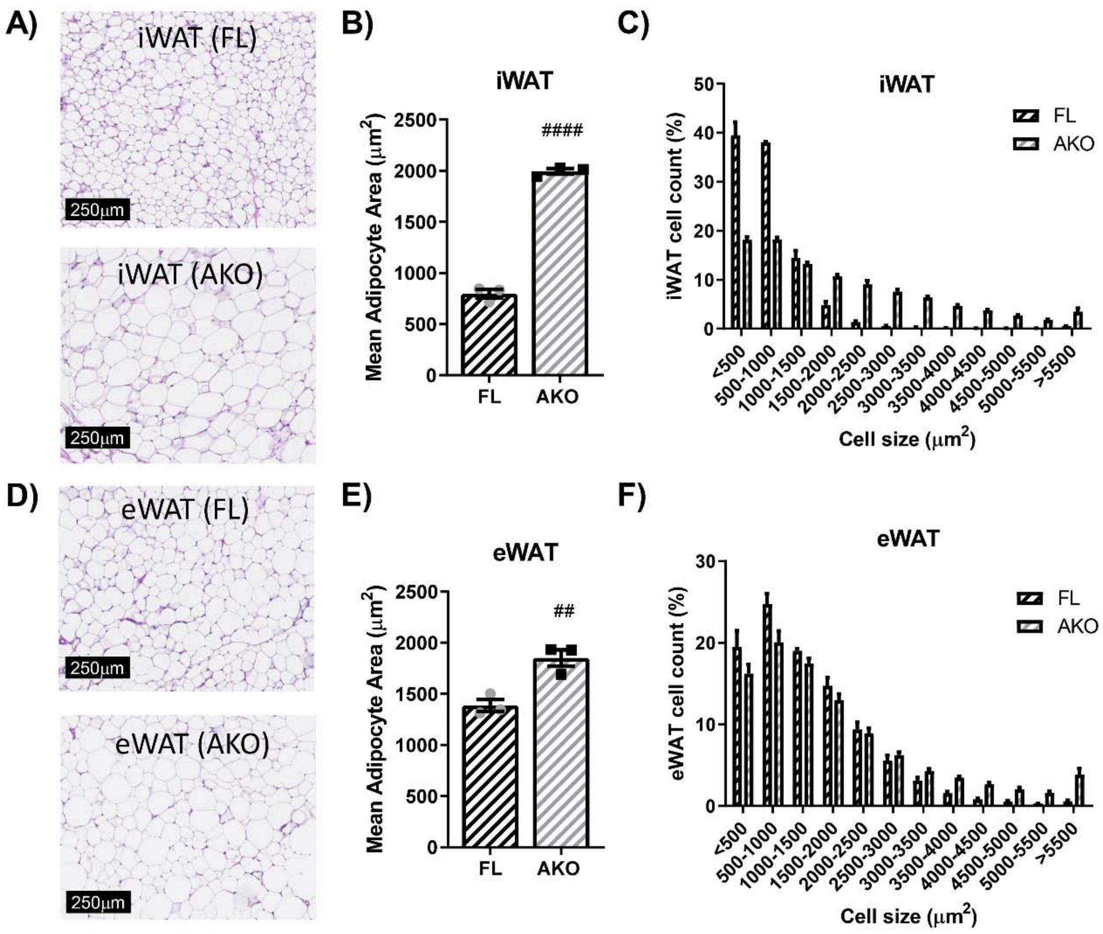


**Figure S3. Male STAT5$^{AKO}$ mice have larger fat cells than floxed control mice in both subcutaneous and visceral adipose tissue depots when fed a chow diet. Related to Figure 3.** Male STAT5$^{AKO}$ (AKO) and floxed (FL) littermate control mice were weaned onto regular chow diet (13% kcal from fat). A and D) Representative images of H&E stained inguinal and epididymal white adipose tissue depots (iWAT and eWAT) collected from five-month-old mice. Quantification of total mean adipocyte area (B and E) and fat cell size distribution (C and F) from H&E-stained images as shown in A and D (n = 3 mice per genotype). Significance in B and E was determined by *t*-test and is denoted as # $p < 0.05$, ## $p < 0.01$, ### $p < 0.001$, or #### $p < 0.0001$ for AKO versus FL comparisons.

**Figure S4. Related to Figure 4 and S5.**

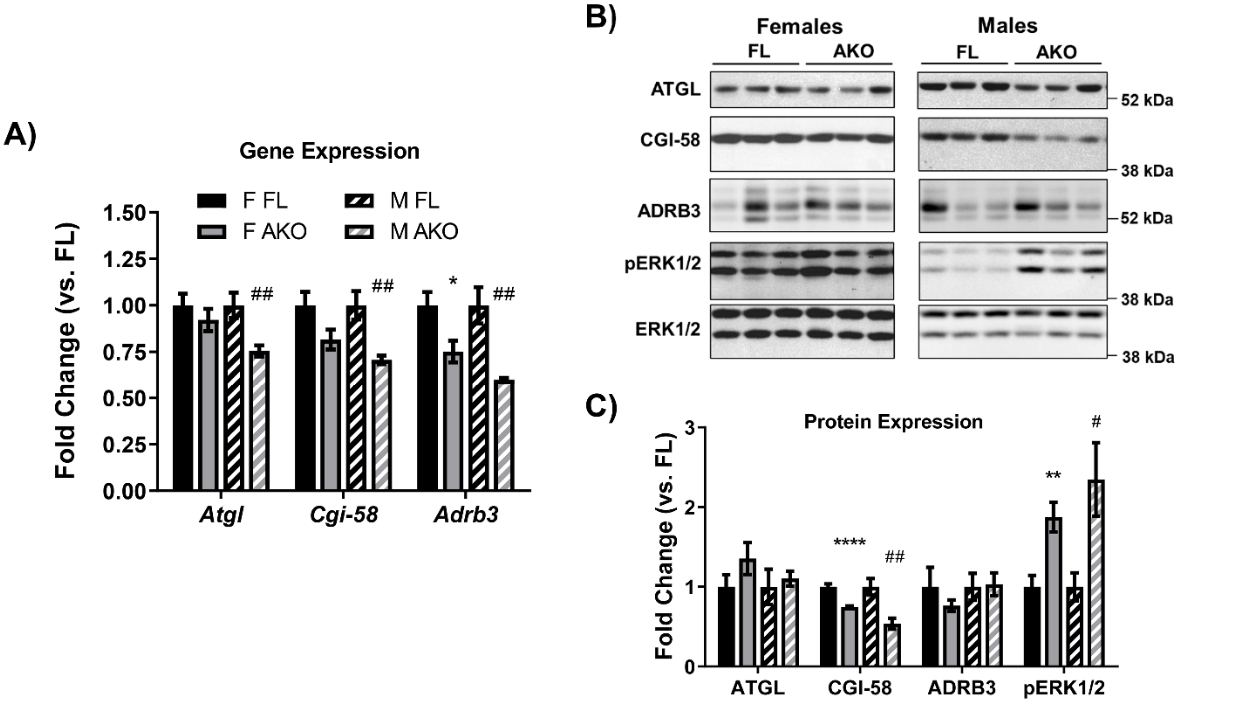


**Figure S4. Effects of loss of adipocyte STAT5 on expression of lipolytic proteins. Related to Figures 4 and S5.** Chow-fed female (F) and male (M) $STAT5^{AKO}$ (AKO) and floxed (FL) littermate control mice were euthanized at 11 weeks of age (non-fasted), and subcutaneous iWAT was collected for gene (A) and protein expression analyses (B and C). A) Gene expression of *Atgl/Pnpla2*, *Cgi-58/Abhd5*, and *Adrb3* was measured by RT-qPCR and normalized against the reference gene *Nono*. B) Protein expression was examined by immunoblotting, and three representative samples per group are shown. C) Band intensities were quantified and normalized against total ERK1/2 expression. Fold change was calculated by dividing the relative gene or protein expression values from each $STAT5^{AKO}$ group by the floxed control group of the same sex for each gene/protein (n = 7 per group). Significance was determined by *t*-test for FL versus AKO comparisons and is denoted as * $p < 0.05$, ** $p < 0.01$, *** $p < 0.001$, or **** $p < 0.0001$ for female comparisons, while significance for male comparisons is denoted as # $p < 0.05$, ## $p < 0.01$, ### $p < 0.001$, or #### $p < 0.0001$.

**Figure S5. Related to Figure 4 and S4.**

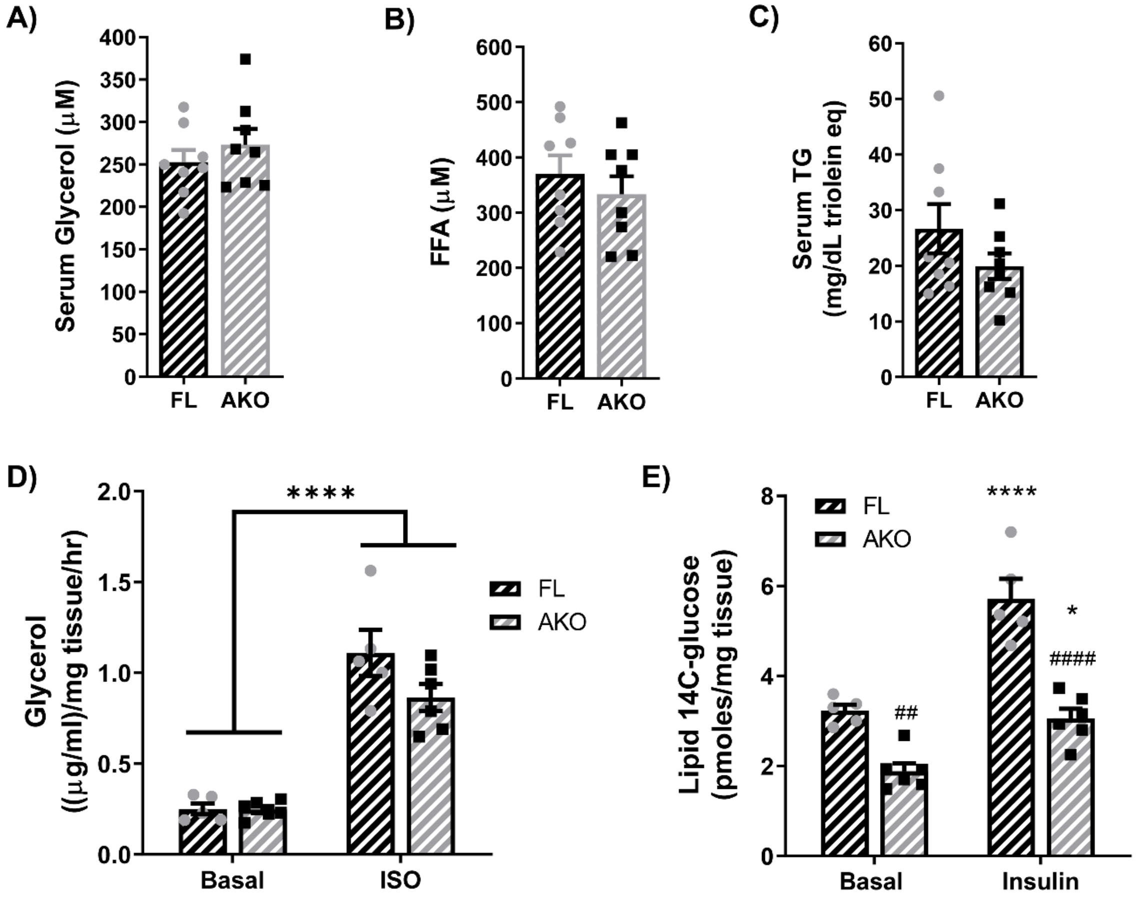


**Figure S5. Male STAT5$^{AKO}$ mice have no differences in markers adipocyte lipolysis compared to floxed control mice, but they do have decreased AT DNL under both basal and insulin stimulated conditions. Related to Figure 4 and S4.** Male STAT5$^{AKO}$ (AKO) and floxed (FL) littermate control mice were fed a defined-composition low fat diet (LFD - 10% kcal from fat; A and B) or regular chow (13% kcal from fat; C – E). A and B) After 1 month on LFD diet and at 3 months of age, serum glycerol and free fatty acid (FFA) levels were measured in blood collected from 4 h-fasted mice. C) Blood was collected from 8-week-old, overnight-fasted mice and serum triglyceride (TG) levels were measured. *Ex vivo* lipolysis (D) and *de novo* lipogenesis (E) assays were performed using gWAT and iWAT explants, respectively, from chow-fed mice (5 months old). D) Glycerol release from gWAT explants (~20mg) into media following a 2 h-incubation period was measured under both basal and isoproterenol (ISO)-stimulated (10μM) conditions. E) Incorporation of $^{14}$C-glucose into total triglycerides was measured by incubating iWAT explants (~50mg) with 4μCi/ml of [$^{14}$C]-U-glucose for 4.5 hours. The triglyceride (neutral lipid) fraction was purified and [$^{14}$C] counts were measured by scintillation counting. For A – C, *t*-tests were used to test for significant between means. D and E) Two-way ANOVA with Tukey's post-hoc multiple comparison analysis was used to test for significance between genotypes and treatments. Significance is denoted as * $p < 0.05$, ** $p < 0.01$, *** $p < 0.001$, or **** $p < 0.0001$ for basal versus ISO or insulin comparisons and # $p < 0.05$, ## $p < 0.01$, ### $p < 0.001$, or #### $p < 0.0001$ for AKO versus FL comparisons.

**Figure S6. Related to Figure 5.**

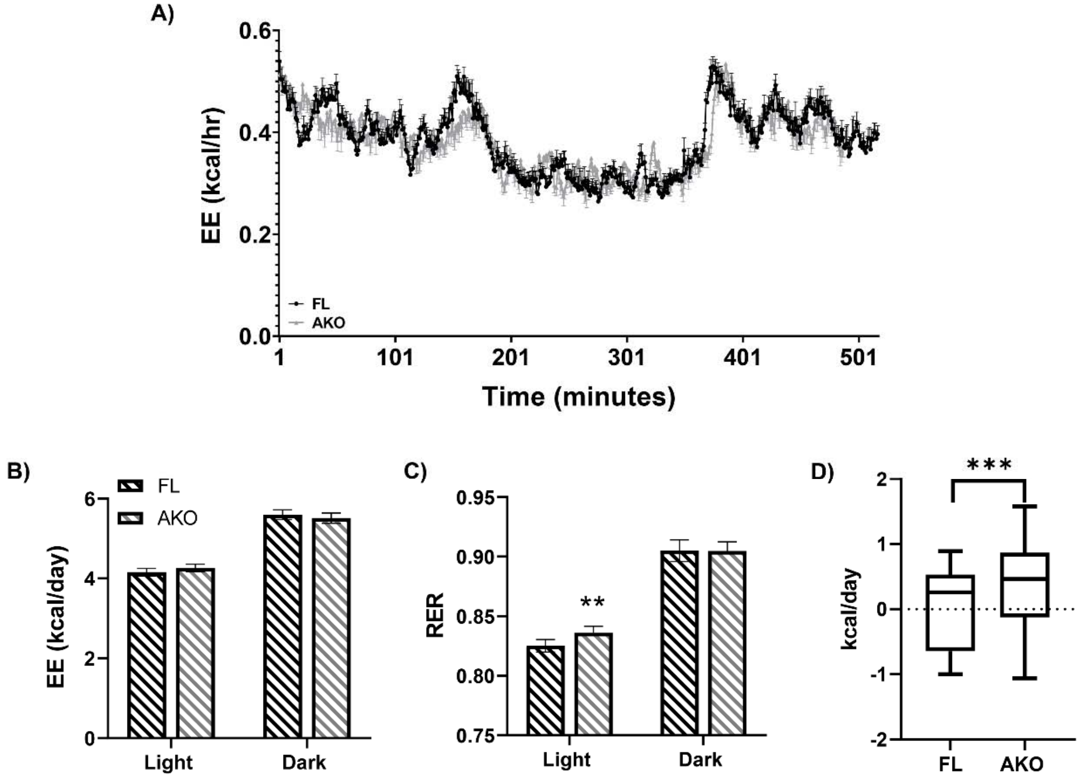


**Figure S6. Loss of adipocyte STAT5 in male mice does not alter energy expenditure but does shift energy balance toward energy storing**. **Related to Figure 5.** Total energy expenditure (EE) and respiratory exchange ratio (RER) for floxed (FL) and STAT5$^{AKO}$ (AKO) male mice (A-C) on a chow diet measured by indirect calorimetry for 7 days (a portion of the longer experiment is shown in A). Energy balance (C) as calculated from the total energy (kcal) consumed minus the total energy expended (kcal). ** denotes $p < 0.01$ and *** $p < 0.001$ (n = 12/group).

**Table S1. Primer sequences for qPCR. Related to STAR Methods.**

| Gene | Primer 1 (5' – 3') | Primer 2 (5' – 3') |
|---|---|---|
| *Cgi-58/Abhd5* | CCCACATCTACATCACACCTT | GAGAGAACATCAGCGTCCATA |
| *Adrb3* | CCACCGCTCAACAGGTTT | CCAGAAGTCCTGCAAAAACG |
| *Cish* | GCTCCTTTCTCCTTCCATCC | CCGCCCAATTTGCTCCA |
| *Atgl/Pnpla2* | GAGCTCATCCAGGCCAAT | CTCATAAAGTGGCAAGTTGTCTG |
| *Stat5a* | ** | ** |
| *Stat5b* | GTTCAACATCAGCAGCAACC | TCAATACTTCCATCACGCCATC |
| *Nono* | CATCATCAGCATCACCACCA | TCTTCAGGTCAATAGTCAAGCC |
| *Cyclophilin a (Ppia)* | TCTTCAGGTCAATAGTCAAGCC | TGCAAACAGCTCGAAGGAGACGC |

** Qiagen RT$^2$ qPCR Primer Assay for Mouse Stat5a – Cat No. PPM04026C-200
