## Supplementary material for "Loss of STAT5 in adipocytes increases subcutaneous fat mass via sex-dependent and depot-specific pathways": Key Resources Table

| REAGENT or RESOURCE | SOURCE | IDENTIFIER |
|---|---|---|
| **Antibodies** | | |
| STAT5A | Santa Cruz Biotechnology<br>Abcam | Cat#: sc-1081 (discontinued); RRID: AB_632448<br>Cat#: ab32043; RRID: AB_778107 |
| STAT5B | R&D | Cat#: AF1584; RRID: AB_2197076 |
| Adiponectin (ADPN) | Thermo Fisher Scientific | Cat#: PA1-054; RRID: AB_325789 |
| ATGL (PNPLA2) | Cell Signaling Technology | Cat#: 2439: RRID: AB_2167953 |
| CGI-58 (ABHD5) | Santa Cruz Biotechnology | Cat#: sc-376931; RRID: AB_2868519 |
| ADRB3 | Santa Cruz Biotechnology | Cat#: sc-515763; RRID: AB_2868520 |
| pERK1/2 (pTEpY) | Promega | Cat#: V8031; RRID: AB_430866 |
| ERK1/2 | Santa Cruz Biotechnology<br>Cell Signaling Technology | Cat#: sc-93 (discontinued); RRID: 631453<br>Cat#: 4695; RRID: AB_390779 |
| **Chemicals, Peptide, and Recombinant Proteins** | | |
| Acrylamide, ProtoGel (30%) | National Diagnostics | EC-890 |
| Collagenase, Type I | Worthington Biochemical | LS004196 |
| Humulin-100<br>(for ITTs) | Eli Lilly | N/A |
| (-)-Isoproterenol (+)-bitartrate | Sigma-Aldrich | I2760 |
| Murine growth hormone | National Hormone and Peptide program (NHPP; Dr. A.F. Parlow) | AFP904 |
| Insulin from bovine pancreas (for lipogenesis assays) | Sigma-Aldrich | I5500 |
| Formalin | Thermo Fisher Scientific | 5725 |
| TRIzol Reagent | Thermo Fisher Scientific | 15596018 |
| **Critical Commercial Assays/Kits** | | |
| Glycerol Assay Kit<br>Glycerol Standard | Sigma-Aldrich | KG0100<br>G7793 |
| NEFA Assay Kit | BioVision<br>Wako diagnostics | K612-100<br>99934691, 99534791, 99134891, 99335191, 27676491 |
| Triglyceride Assay Kit | Sigma-Aldrich | TR0100 |
| Mouse Insulin ELISA | Crystal Chem | 90080 |
| Mouse IGF-1 ELISA | Crystal Chem | 80574 |
| Mouse GH ELISA | Millipore Sigma | EZRMGH-45K |
| SYBR® Premix Ex Taq (Tli RNase H Plus), Rox Plus | Takara Bio USA | RR42LR |
| Quant-Seq 3' mRNA-Seq Library Prep Kit FWD for Illumina | Lexogen | 015.2X96 |
| **Deposited Data** | | |
| Raw and analyzed RNA-seq data | This paper | GEO: GSE113939 |
| **Experimental Models: Organisms/Strains** | | |
| B6.FVB-Tg(Adiopq-cre)1Evdr/J | The Jackson Laboratory | 028020 |
| $STAT5^{fl/fl}$ | (Cui et al., 2004) | N/A |
| $STAT5^{AKO}$ | This paper | N/A |

| | | |
|---|---|---|
| Oligonucleotides | | |
| Primers for qPCR | Table S1 | N/A |
| Software and Algorithms | | |
| Graph Pad Prism 6 & 8.4 | GraphPad Software | N/A |
| Image Studio Lite Ver 5.2 | LI-COR Biosciences | N/A |
| NanoZoomer Digital Pathology; NDP.view 2.7.52 | Hamamatsu | N/A |
| QuantSeq pipeline V1.8.8 on Bluebee platform | https://www.lexogen.com/store/quantseq-data-analysis-bluebee-platform/ | N/A |
| Other | | |
| Chow Diet | LabDiet® | 5001 and 5015 |
| Low Fat Diet (LFD), no sucrose | Research Diets | D12450K |
| High Fat Diet (HFD) | Research Diets | D12492 |
| High Fat High Sucrose (HFHS) Diet | Research Diets | D08112601 |
| Sarstedt Capillary blood collection tube | Fisher Scientific | NC9059691 |
| NMR Machine | Bruker | Minispec LF110 |
| 7900HT qPCR Machine | Applied Biosystems | N/A |
| Metabolic Chambers | Sable Systems International | Promethion |
| NextSeq 500 Sequencer | Illumina | N/A |
